# Structure of the Human Signal Peptidase Complex Reveals the Determinants for Signal Peptide Cleavage

**DOI:** 10.1101/2020.11.11.378711

**Authors:** A. Manuel Liaci, Barbara Steigenberger, Sem Tamara, Paulo Cesar Telles de Souza, Mariska Gröllers-Mulderij, Patrick Ogrissek, Siewert J. Marrink, Richard A. Scheltema, Friedrich Förster

**Author notes:** Correspondence to l.

## Abstract

The signal peptidase complex (SPC) is an essential membrane complex in the endoplasmic reticulum (ER), where it removes signal peptides (SPs) from a large variety of secretory pre-proteins with exquisite specificity. Although the determinants of this process have been established empirically, the molecular details of SP recognition and removal remain elusive. Here, we show that the human SPC exists in two functional paralogs with distinct proteolytic subunits. We determined the atomic structures of both paralogs using electron cryo-microscopy and structural proteomics. The active site is formed by a catalytic triad and abuts the ER membrane, where a transmembrane window collectively formed by all subunits locally thins the bilayer. This unique architecture generates specificity for thousands of SPs based on the length of their hydrophobic segments.

## Introduction

**M**any secretory pathway proteins are targeted to the endoplasmic reticulum (ER) *via* a short N-terminal transmembrane helix called signal peptide (SP) (1). Nascent SPs emerge from the ribosome and target the ribosome-nascent-chain complex to the ER membrane, where it is inserted into the protein-conducting channel Sec61. For many proteins (approximately 5,000 different physiological protein substrates in humans (2)), the signal peptidase complex (SPC) cleaves off the SPs from their non-functional pre-forms. The SPC also facilitates the maturation of many viral proteins, including pre-proteins from most flaviviruses (e.g. Zika, Dengue, and Hepatitis C virus), HIV, and SARS coronavirus (3–7).

The human SPC comprises the accessory proteins SPC12 (SPCS1), SPC22/23 (SPCS3), SPC25 (SPCS2) and the two proteolytic subunits SEC11A (SPC18) and SEC11C (SPC21) (Fig. 1A) (8). It is currently unclear whether both proteolytic subunits occur in the same complex or form distinct SPC paralogs (9). Both SEC11A and SEC11C have low but significant sequence similarity to bacterial signal peptidases (SPases) (10), which are monomeric and characterized by a Lys-Ser catalytic dyad (11, 12). In contrast, eukaryotic SPCs have the active site lysine replaced by a histidine, and might either function through a catalytic His-Ser dyad or Asp-His-Ser triad (13), leading to the functional distinction of prokaryotic P-type SPases and eukaryotic ER-type SPases (14).

**Figure 1.**
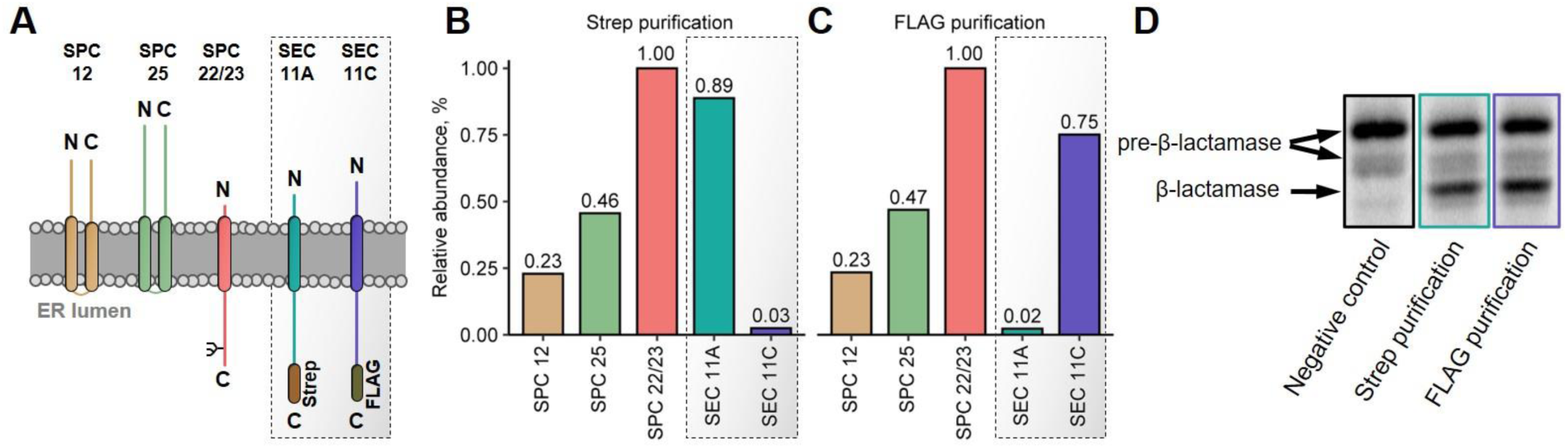
SPC exists in two paralogs. (**A**) Overview of the SPC subunits. The non-proteolytic subunits SPC12 (yellow), SPC25 (green), and SPC22/23 (red) were co-expressed with SEC11A-Strep (teal) and SEC11C-FLAG (purple). (**B-C**) Top-down MS quantification of the subunits after Strep (**B**) or FLAG (**C**) affinity purification. Abundance normalized to SPC22/23. (**D**) Pre-β-lactamase *in vitro* cleavage assay. Negative control = no SPC added.

The SPC is highly selective for SPs, but the molecular mechanism of SP recognition is largely unexplored. Consequently, SPs are typically predicted using empirical features (15). SPs are characterized by three distinct regions: (*i*) an often positively charged, unfolded n-region, (*ii*) a hydrophobic, alpha-helical h-region and (*iii*) a polar c-region, which contains the scissile bond (16). The 1-5 residue n-region determines the orientation of the SP in the conducting channel Sec61 and hence membrane protein topology (17). The h-region of SPs is invariably hydrophobic and with 7-15 amino acids notably shorter than regular TM helix segments (18). The c-region is 3-7 amino acids long and contains two crucial positions relative to the scissile bond (−1 and -3) that need to be occupied by small, non-charged residues.

We reconstituted the human SPC and analyzed it by cryo-EM single particle analysis and structural proteomics-driven mass spectrometry (MS) in order to elucidate its precise stoichiometry, structure, and the mechanism of SP recognition and cleavage.

## Results & Discussion

### Human SPC exists in two paralogs

SPC12, SPC22/23, and SPC25 can be found in essentially all eukaryotes, suggesting they have evolved at the advent of eukaryotic life (**Fig. S1**). In most eukaryotic organisms, the SPC only consists of these three subunits and one copy of SEC11. In animals, a duplication event of SEC11 occurred approximately 400 mio. years ago. SEC11A and SEC11C remained closely related throughout evolution, with ∼80% sequence identity in humans. Both genes can individually substitute yeast SEC11 and even some bacterial SPases functionally (19).

To test whether SEC11A and SEC11C are part of two distinct paralogous SPCs, we co-expressed the three accessory subunits SPC12, SPC25 and SPC22/23 with Strep-tagged SEC11A and FLAG-tagged SEC11C in HEK 293 cells and purified the complexes by either Strep or FLAG affinity chromatography from the same batch of cells (Fig. 1, **S2**). In both cases, we recovered near-stoichiometric amounts of the accessory subunits and the respective tagged SEC11 variant, while the other variant was 30-40 times less abundant as determined by top-down mass spectrometry (Fig. 1B-C). Both isolates were able to process pre-β-lactamase *in vitro* with similar efficiencies (Fig. 1D). We conclude that in humans, and likely in other eukaryotes with two SEC11 paralogs, two functional hetero-tetrameric SPC paralogs exist formed by SPC12, SPC22/23, SPC25 and either SEC11A or SEC11C. In the following, we refer to the two paralogous complexes as SPC-A and SPC-C, respectively.

### SPC architecture and topology

We determined the structures of both human paralogs, solubilized in amphipol PMAL-C8, using single particle cryo-EM to an overall resolution of approximately 4.9 Å (Fig. 2A-B, **S3-S5, Table S1, Movie S1**). The low protein mass of the hetero-tetrameric complex (84 kDa, 17 of which are unordered) and the structural variations of the micelle likely limited particle alignment accuracy and attainable resolution (20). Initial atomic models of the subunits generated by trRosetta (21) yielded excellent fits to the two cryo-EM densities (**Fig. S5**). Using these initial models and the EM maps, we could build atomic models of both SPCs that explain all of the observed density (Fig. 2C, **S5**). The models agree with the previously determined transmembrane topologies of the subunits (22, 23) as well as predictions of transmembrane helices, secondary structure and disordered segments, and atomistic molecular dynamics (MD) simulations (**Fig. S6**). When mapping the distance restraints obtained by XL-MS onto the atomic model, we found that ∼80% of the cross-links range within the maximum allowed distance of the PhoX crosslinker (24) (**Fig. S7**).

**Figure 2.**
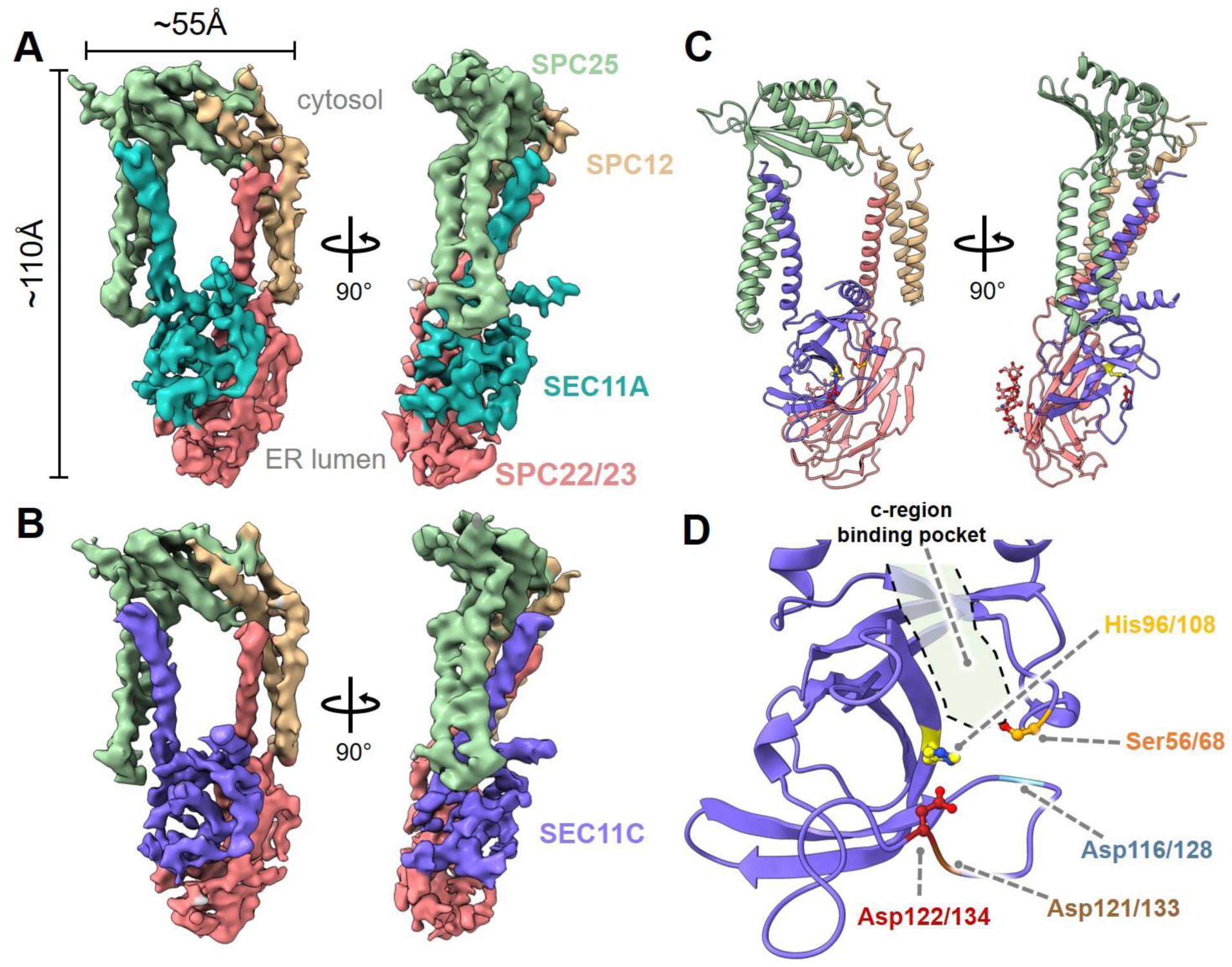
Structure and SP binding pocket of the human SPC. (**A**) EM map of the SPC-A complex, with density for SEC11A in teal, SPC22/23 in red, SPC25 in green, SPC12 in yellow. (**B**) EM map of SPC-C. SEC11C is colored purple. (**C**) Atomic model of SPC-C. (**D**) Conserved SPase I fold of SEC11C. The c-region binding pocket and (candidate) catalytic residues are highlighted.

In the structures, SEC11A or SEC11C, respectively, interacts with SPC22/23 to form a globular luminal body consisting solely of beta sheets (Fig. 2C). The transmembrane domains of SPC25 and SEC11A/C on one side and SPC12 and SPC22/23 on the opposite side form two distinct three-helix bundles, which frame a characteristic ∼15 Å wide lipid-filled transmembrane window (‘TM window’) in the membrane. The cytosolic portion of the complex is formed mainly by SPC12 and SPC25. Together, these two subunits form a clamp-like structure that orients the transmembrane segments of SEC11A/C and SPC22/23 (Fig. 2C).

Consistent with this architecture, native MS shows detectable SEC11A/C-SPC22/23 sub-complexes, which dissociate at comparable activation energies (**Suppl. Text, Fig. S8**). SPC25 and SPC12 were detected only in a free form, likely because removal of the amphipol affected their binding interfaces. The cryo-EM density explains ∼80% of the SPC residues, whereby most of the unresolved residues are mapping to the N-termini of SPC12 and SPC25 (**Fig. S6**). We detected the bulk of the unmapped N- and C-terminal regions of SEC11A/C, SPC25 and SPC12 by shotgun and top-down mass spectrometry, which confirms that they are structurally flexible rather than proteolytically removed (**Fig. S9**). The terminal stretches of the SPC harbor different PTMs, such as phosphorylation (SPC12), N-terminal acetylation (SPC25) and partial N-terminal truncation (all subunits except for SPC22/23 and SEC11A) (**Fig. S9**).

### Characterization of the accessory subunits

The luminal domain of SPC22/23 forms an extended beta sandwich with a fold similar to that of the histone chaperone ASF1 (25), which embraces the catalytic core of SEC11 (Fig. 2C). This arrangement suggests that SPC22/23 helps to stabilize and position the active center close to the luminal membrane surface, which explains why it is required for catalytic function of the SPC (26–28). In our sample derived from HEK 293 cells, we found 98% of SPC22/23 proteins to be N-glycosylated at Asp141, which is considerably higher than reported for dog pancreatic microsomes (8). Top-down mass spectrometry identified this glycan as a homogeneous biantennary mannose-type structure (**Fig. S9**). It is also partially resolved in the EM map and projects towards the membrane.

The subunits SPC12 and SPC25 are not essential for catalytic activity (29), but deletion of SPC25 in yeast results in a two-fold reduction of *in vitro* SPase activity (30). In our structures, SPC25 accounts for most of the ordered density in the cytosolic portion of the SPC. The protein adopts a novel alpha-beta sandwich fold, which is interspersed by the two transmembrane helices that interact with SEC11A/C. The N-terminal 50 amino acids of SPC25 are missing from the density but detected by various MS approaches. In accordance with previous reports (31), the removed starting methionine at the N-terminus of SPC25 is replaced by an N-acetylation. In addition, a subset of SPC25 molecules is N-terminally processed (**Fig. S9**).

SPC12 is the only subunit that does not directly interact with SEC11 (Fig. 2C), explaining why it is least important for catalytic activity (30, 32). The cytosolic termini of SPC12 are largely flexible (residues 1-65 and 152-169), and only its membrane-proximal parts constantly interact with SPC25 as supported by XL-MS data (**Fig. S7**). SPC12 exhibits minor N-terminal processing as revealed by top-down mass spectrometry along with a low-stoichiometric phosphorylation which is likely located on the cytosolic portion of the complex (**Fig. S9**).

### Characterization of SEC11 and the SP c-region binding pocket

The luminal SEC11A/C portion adopts an SPase I fold that aligns well with the catalytic core domain of *E*.*coli* SPase I (12) (Fig. 2D, **S10**). Due to the low homology between P- and ER-type SPases, the interspersed conserved sequence stretches are commonly referred to as boxes A-E (14) (**Fig. S10A-B**). The catalytic residues Ser56/68 (in box B, numbered as in SEC11A/C, respectively) and His96/108 (in box D) are located at highly similar positions as the SPase I Ser-Lys dyad. In contrast to the P-type SPases, it has previously been suggested that ER-type SPases might function through a catalytic Ser-His-Asp triad because SEC11 has three conserved aspartic acid residues, Asp116/128, Asp121/133 and Asp122/134 (all box E), that might complete the active center (13) (Fig. 2D, **S11**). Our structures show that all three aspartic acid residues are located proximal to the binding pocket, with Asp122/134 best positioned to complete the triad. The models suggest that Asp116/128 points towards the protein core and engages in a salt bridge with the equally conserved Arg97/109, analogous to structures of P-type SPases (12, 33). Since the map resolution is insufficient to model side chains reliably, we mutated all three candidate aspartic acids and tested how they affect catalytic activity and protein stability *in vitro* (**Fig. S12**). Mutating Asp122/134 had only a moderate effect on protein stability while completely abolishing catalytic activity. As expected, mutating Asp116/128 had a more severe effect on protein stability, but retained catalytic activity to a reasonable extent, while mutating Asp121/133 had little effect on both stability and activity. We thus conclude that human SEC11A/C indeed function *via* a catalytic triad consisting of Ser56/68, His96/108, and Asp122/134.

To model the c-region of SPs in the SPC, we superposed SEC11A/C and *E*.*coli* SPase I in complex with the lipopeptide inhibitor arylomycin (34) as a template for the c-region (**Fig. S10C-G**). As in its bacterial counterpart, the catalytic residues of SEC11A/C reside at the end of a shallow, hydrophobic groove that is lined by a β-strand formed by box D residues. In the bacterial enzyme (and most other proteases), the c-region of the SP is forced into a β-strand conformation (35, 36). The substrate side chains at the -1 and -3 positions point towards shallow hydrophobic pockets that can only accommodate small hydrophobic residues (35). The same principle likely applies to SEC11A/C and provides an explanation for the empirically established c-region consensus motif and the interchangeability of bacterial and eukaryotic SP c-regions (**Fig. S10F-G**).

Both SEC11A and SEC11C possess a single, N-terminal TM helix, which was not resolved in structures of bacterial homologs (12). Near the C-terminus, SEC11A/C have a striking amphiphilic helical segment at the interface between membrane and ER lumen that we termed the ‘bowsprit helix’ because it prominently projects from the binding pocket (Fig. 3B,E). The N- and C-terminal stretches of SEC11A/C, which harbor most of the single amino acid variations, are flexible in our structures.

**Figure 3.**
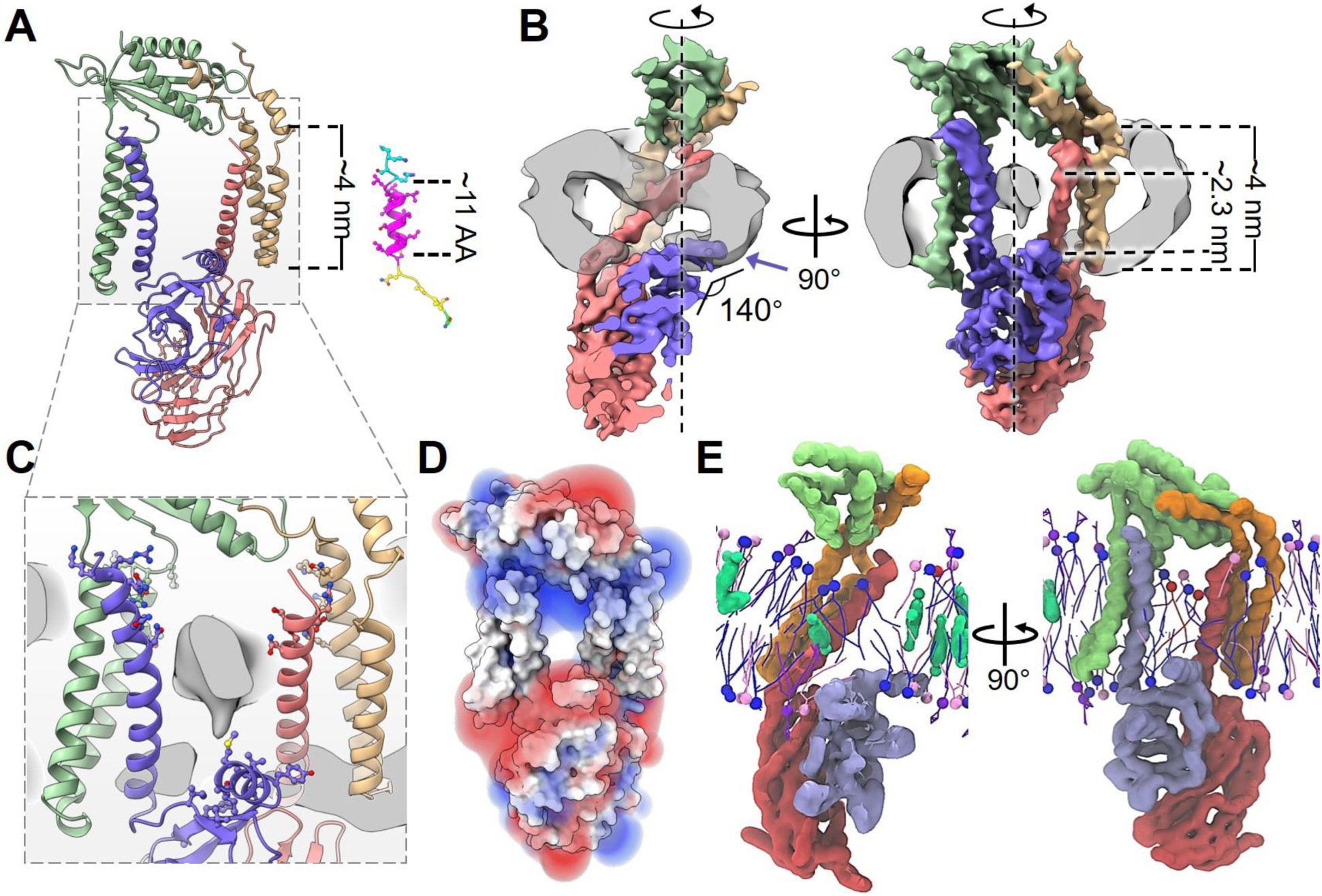
The SPC locally thins the membrane. (**A**) SP h-regions are short compared to the SPC transmembrane helices. The SP of bovine pre-prolactin is shown (cyan = n-region; magenta = h-region; yellow = c-region). (**B**) Slices through a micelle-containing SPC-C map demonstrate local membrane thinning. Dimensions of the membrane inside and outside the TM window are given. Bowsprit helix indicated with a purple arrow. SEC11C presses against the membrane from the lumen. (**C**) Polar residues lining the inside of the TM window. The hydrophobic ridge of SEC11C is shown on the luminal side. (**D**) Electrostatic fields on both sides of the TM window (blue = positive; red = negative). (**E**) Coarse-grained MD simulation in a complex ER bilayer showing the lipid distribution within the TM window (blue = POPC; red = POPS; pink = POPE; purple = PI(3,4)P2; green = cholesterol).

### Membrane shaping by the SPC allows SP recognition based on the h-region

On the inside of the lipid-filled ‘TM window’, the diameter of the amphipol micelle is reduced to approximately 23 Å compared to 35-40 Å in the exterior (Fig. 3). This thinning of the micelle also occurs when the SPC is solubilized in digitonin (**Fig. S13**). Coarse grained and atomistic MD simulations confirm that the thinning is also present in simple and complex lipid membranes (Fig. 3E and **Fig. S14**). For instance, in a complex ER-like membrane, we observed an average thinning of 26%, with fluctuations between 15-46%. As a consequence, the TM window seems enriched of lipids which usually form thin membranes (37) - especially unsaturated phosphatidylcholine lipids - which spread their acyl chains to squeeze into the window (**Fig. S14**). Thus, the SPC induces a local thinning of the lipid membrane reminiscent of protein insertase and translocation complexes such as YidC, EMC, and the Hrd1-ERAD complex (38).

The SPC structure suggests that several factors synergistically induce membrane thinning: (i) on the cytosolic face of the SPC, the sides of the three-helix bundles that frame the TM window have notably shorter hydrophobic cores than those facing the surrounding membrane, and they are notably positively charged at their cytosolic ends (Fig. 3C,D). (ii) On the luminal face of the window, a range of membrane-proximal residues contribute to a negative charge (Fig. 3D), while (iii) SEC11A/C forms a hydrophobic ridge that presses tightly against the membrane and partially inserts itself into the hydrophobic environment (Fig. 3B,C). The SEC11 bowsprit helix is prominently positioned on the micelle surface, suggesting that it contributes to shaping the membrane surrounding the binding pocket (Fig. 3B,E).

The entire c-region of SPs measures five to six amino acids on average, which fits the distance from the active site to the thinnest point of the amphipol located right above the SP binding groove (**Fig. S10G**). At this position, the micelle diameter (∼23 Å) coincides with the length of a typical h-region (∼11 amino acids) (Fig. 3A,B). One of the main determinants distinguishing SPs from other TM helices is its short h-region. Hence, the membrane thinning in the SPC window may promote preferred accommodation of short h-regions and thus be key for SPC specificity (Fig. 4A, **Movie S2**). In this context, the enrichment of phosphatidylcholine within the TM window indicated by our MD simulations also explains why relipidation of the SPC with phosphatidylcholine is required to restore the catalytic activity of the SPC in some detergent systems (8, 39, 40). The observation that the catalytic domain of bacterial SPase I has higher affinity for lipid monolayers than for bilayers suggests that this mechanism is conserved (41).

**Figure 4.**
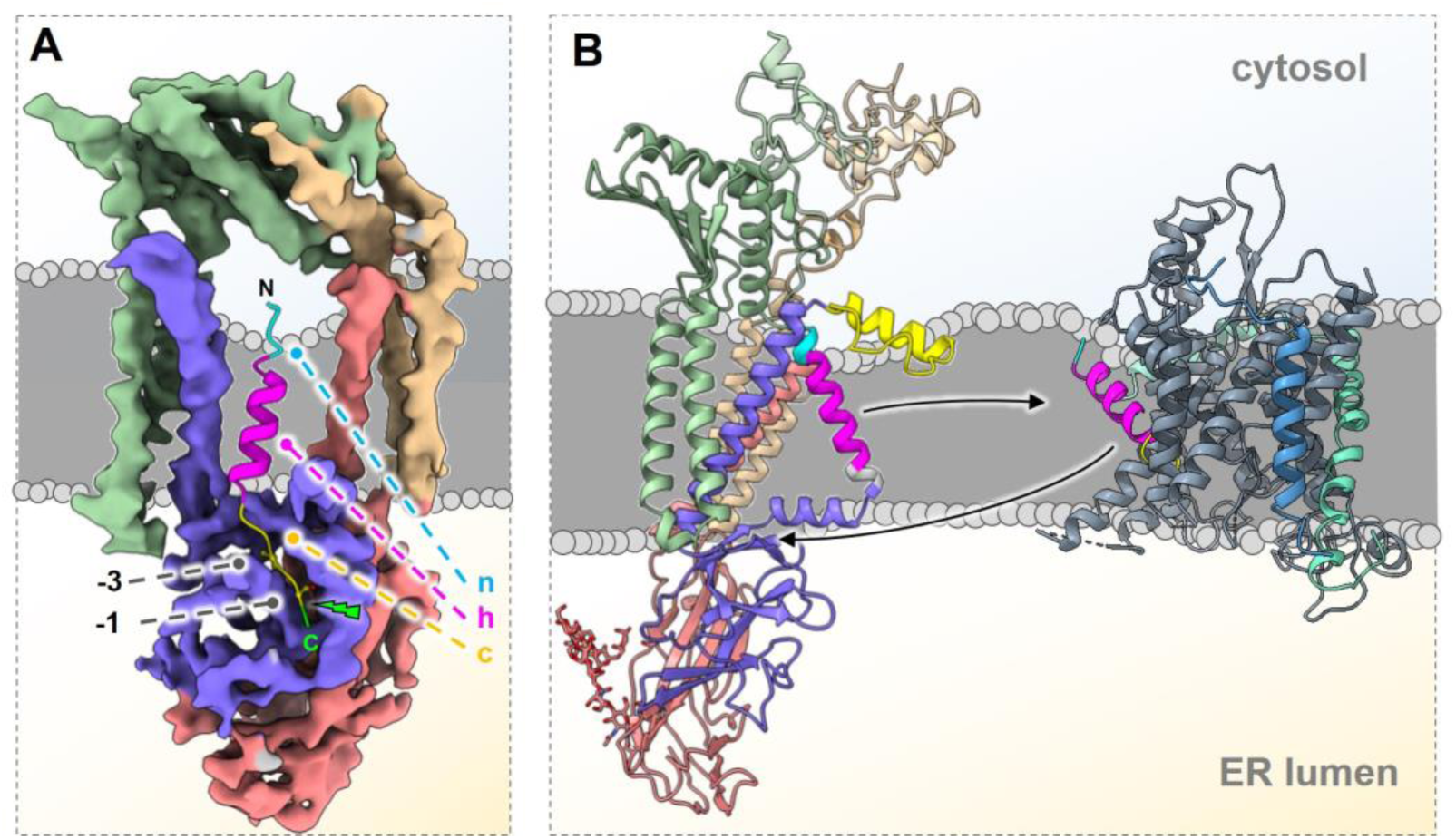
Models for SP engagement by the SPC. (**A**) Proposed model of SP engagement. The SP of bovine pre- prolactin has been modelled into the SPC-C binding pocket. The SP (colored as in Fig. 3) is recognized based on h-region length and shape complementarity in the c-region. The scissile bond and start of mature sequence are highlighted in green. (**B**) SP hand-over model for Sec61 interaction. Sec61 complexed with bovine pre-prolactin (PDB ID 3JC2) shown as ribbons (Sec61α = grey; Sec61β = blue; Sec61γ = green). The pseudo-SP helix might displace the SP from Sec61 by binding to the lateral gate. The displaced SP might be guided into the SPC active center by the gradient of the membrane thinning and the N-terminal amphipathic helix and bowsprit helix of SEC11.

### Differences between the two paralogs

It is currently unclear whether the two SPC paralogs play distinct roles in substrate processing. Substitution of SEC11 by either SEC11A or SEC11C in yeast indicated only subtle functional differences for a small set of substrates (19), and processing of flaviviral pre-proteins was largely unaffected by either SEC11A or SEC11C knockout in HEK cells (7). In an attempt to get further clues on substrate specificity of SPC-A and SPC-C we determined the relative abundance of SEC11A and SEC11C in a number of common cancer cell lines (**Fig. S15A**). In all these cell lines, SEC11A is highly expressed, while the level of SEC11C is below the detection limit. Nevertheless, SEC11C is reported to be ubiquitously expressed in many tissues at similar levels as SEC11A (42).

The maps for SPC-A and SPC-C are virtually indistinguishable at the current resolution (**Fig. S15B-E, Movie S1**). The residues mapping to the N-terminal SEC11 transmembrane helix and the SP binding groove are completely conserved, while the sequence variations (located mostly on surface-exposed loops on the periphery of the SPase domain) do not result in significant structural differences (**Fig. S15F-H**). However, there are substantial sequence differences in the flexible N- and particularly the C-terminal stretches of SEC11A and SEC11C (**Fig. S15G-H**). The cytosolic N-termini of both SEC11A and SEC11C are predicted to form short, amphiphilic helical segments (**Fig. S16**). The N-terminal segment of SEC11C is 12 residues longer than SEC11A. These residues are conserved among mammals (**Fig. S11**), despite being predicted to be unstructured (**Fig. S6**). Top-down MS reveals significant sequence processing for SEC11C in this area with removal of up to nine N-terminal residues, while SEC11A appears as a single unprocessed and unmodified proteoform (**Fig. S9**).

At their C-termini, both SEC11 paralogs are predicted to possess a helix that connects with the bowsprit helix through a conserved proline residue. This ‘helix breaker’ splits the two segments at the interface between lumen and ER membrane (**Fig. S16**). Interestingly, the primary sequence of this C-terminal helix has a hydrophobic stretch of 13 amino acids and resembles an inverted SP, analogous to type-III signal anchors (**Fig. S16C-D**). We therefore named this segment the ‘pseudo-SP helix’. Given that GFP fused to the C-terminus of SEC11C is found on the cytosolic side of digitonin-solubilized particles (**Fig. S13C**) and crosslinks only to cytosolic portions of the SPC (**Fig. S7B**), we employed atomistic MD simulations to test whether the pseudo-SP helix could span a lipid membrane (**Fig. S6C**). The data shows that the pseudo-SP helix, which is not resolved in the EM map, can indeed be stably accommodated as a transmembrane helix in the thinned membrane environment of the SPC, much like actual SPs. This helix might directly interact with the h-region of SPs and confer some sort of selectivity for different subgroups of SPs.

### Functional model for co-translational SP cleavage

At the ER, the SP of the nascent peptide is initially accommodated in the lateral gate of the Sec61 protein-conducting channel (43–45), which forms the core of the ER translocon complex. Given that SPs are stoichiometrically bound to the native ER translocon (46), the association of the SP to Sec61 appears long lived in the cell. For cleavage, the SP needs to transfer from Sec61 to the SPC, which co-purifies with the ER translocon (47). However, the SPC is not resolved as part of native ribosome-translocon complexes by cryo-electron tomography (43, 48), indicating that the SPC likely associates with the ER translocon in a structurally flexible manner (49). It was demonstrated previously that SPC25 and Sec61 can be cross-linked at their flexible parts in both yeast and mammals (30, 50). Topologically, the existence of these cross-links suggests that the active center of the SPC faces Sec61, with the bowsprit and pseudo-SP helices pointing towards the lateral gate of Sec61. Given the similarity between the pseudo-SP helix and SPs, we speculate that the pseudo-SP helix is involved in transfer of the SP from Sec61 to the SPC, for example by transient association with the Sec61 lateral gate triggering co-translational release of the SP into the locally thinned membrane of the SP window (Fig. 4B).

In summary, this study provides a basic understanding of how the SPs of thousands of substrates are recognized and cleaved by the SPC. It also serves as a foundation for the characterization of interactions between the SPC and viral preproteins and their pharmacological interference, as well as the development for SPase-targeting antibiotics that do not affect the human SPC.

## Methods

### Design and cloning of expression constructs

All subunits of SPC-A (SPC12,25,22/23, and SEC11A) and SPC-C (SPC12,25,22/23, and SEC11C), respectively, were expressed from a single pUPE-2961 vector (U-Protein Express BV) in one large ORF (**Fig. S2, S17**). The individual subunits were separated by picornaviral 2A modules. Codon-optimized DNA constructs based on UniProtKB entries Q9Y6A9-1 (SPC12), Q15005 (SPC25), P61009 (SPC22/23), P67812 (SEC11A), and Q9BY50 (SEC11C) were synthesized by Twist Bioscience and cloned into the vector backbone by Gibson assembly (New England Biolabs) in the sequence SPC25-[E2A]-SPC12-[P2A]-SPC22/23-[T2A]-SEC11A (SPC-A) or SPC25-[E2A]-SPC12-[P2A]-SPC22/23-[T2A]-SEC11C (SPC-C). The respective SEC11 subunit was C-terminally tagged with a TEV-cleavable eGFP-TwinStrep-HA tag.

For composition analysis, proteolytic subunits were removed from the expression constructs using blunt end deletion following a standard Q5 mutagenesis workflow (New England Biolabs). Additionally, SEC11A and SEC11C were separately cloned into pUPE-2961 by Gibson assembly. A C-terminal Strep tag was added to SEC11A and a C-terminal FLAG tag was added to SEC11C by Q5 mutagenesis, and *vice versa*. Similarly, point mutants were generated from the parental constructs by Q5 mutagenesis. All constructs were evaluated by sequencing.

### Protein Expression and Purification

#### Expression

All SPC constructs were transiently expressed for ∼48h in suspension HEK 293-E^+^ cells by U-Protein Express BV (Utrecht, the Netherlands) using 0.5 mg vector DNA per L cell culture. The final cell densities ranged between 1-2 million cells per mL. All subsequent steps were performed at 4 °C unless stated otherwise. Cells were pelleted by centrifugation at 500 g and washed three times with ice-cold PBS to remove biotin from the expression medium. The resulting cell pellets were flash-frozen in 0.5 L aliquots and stored at -80 °C until further use. For composition analysis, cells from 1 L culture were co-transfected with 0.333 mg of the vector containing the accessory subunits SPC25, SPC12, and SPC22/23 as well as 83.3 μg of SEC11A-Strep and SEC11C-FLAG, respectively.

#### For functional assays: Purification in digitonin

Cell pellets from 0.2-0.5 L culture medium were thawed in 35 mL lysis buffer per L cell culture (50 mM HEPES pH 7.8 100 mM NaCl 5 mM EDTA, 1 mM DTT, 10% (v/v) glycerol, 0.7 μg/mL DNase I, 1% (w/v) digitonin) and incubated 1.5 h at 4 °C in a rotating wheel. Since the SPC is resistant to common protease inhibitors (51), one cOmplete inhibitor tablet (Roche, containing EDTA) was added during cell lysis. Samples were cleared by ultracentrifugation at 100,000 g for 30 min in a fixed-angle rotor. The resulting supernatant was immobilized twice on 5 mL pre-equilibrated Streptactin XT high capacity beads (IBA) in a gravity flow column, and the immobilized sample was washed with 20 column values (CV) wash buffer (20 mM HEPES pH 7.8, 85 mM NaCl, 1 mM EDTA, 1 mM DTT, 10% (v/v) glycerol, 0.1% (w/v) digitonin). Retained SPC was eluted with 5-10 CV elution buffer (100 mM HEPES pH 7.8, 85 mM NaCl, 1 mM EDTA, 1 mM DTT, 10% (v/v) glycerol, 0.09% (w/v) digitonin, 50 mM biotin). These preparations were used for XL-MS and to analyze the SPC point mutants (**Fig. S17**). For the latter, samples were buffer-exchanged (1:400) into wash buffer and concentrated to 1.0 mg/mL.

A similar protocol was used for composition analysis, except that DTT was omitted from all buffers until affinity chromatography was completed. 300 mL of centrifuged cell lysate were split into two batches and immobilized four times on two gravity flow columns - one containing 0.5 mL Streptactin XT high capacity beads (IBA) and one containing 0.5 mL anti-FLAG M2 affinity resin (Sigma-Aldrich) (**Fig. S2**). After each round of immobilization, the flow through from each column was re-applied to the other column. Both columns were washed with 40 CV of wash buffer without DTT before elution at room temperature with 10x 1 CV of elution buffer without DTT. For the FLAG resin, 50 mM biotin were replaced with 200 μg/mL 3x FLAG peptide (Sigma-Aldrich). Samples were buffer-exchanged into wash buffer containing DTT (dilution factor 1:400) and concentrated to 1 mg/mL.

#### For cryo-EM: Purification in PMAL-C8

Cell pellets were thawed in 35 mL EM lysis buffer (50 mM HEPES pH 7.8, 100 mM NaCl 5 mM EDTA, 1 mM DTT, 10% (v/v) glycerol, 1 large Roche cOmplete inhibitor tablet (containing EDTA), 0.7 μg/mL DNAse I, 1% (w/v) DDM, and 0.2% (w/v) CHS) per L cell culture and incubated 1.5 h at 4 °C in a rotating wheel. Samples were cleared by ultracentrifugation at 100,000 g for 30 min in a fixed-angle rotor. The resulting supernatant was immobilized twice on 5 mL Streptactin XT high capacity beads in a gravity flow column, and the immobilized sample was washed with 20 CV EM wash buffer (20 mM HEPES pH 7.8, 85 mM NaCl, 1 mM EDTA, 1 mM DTT, 10% (v/v) glycerol, 0.0174% (w/v) DDM, 0.00348% (w/v) CHS). Retained SPC was eluted with 5-10 CV EM elution buffer (100 mM HEPES pH 7.8, 85 mM NaCl, 1 mM EDTA, 1 mM DTT, 10% (v/v) glycerol, 0.0174% (w/v) DDM, 0.00348% (w/v) CHS, 50 mM biotin). The eluate was concentrated to 1 mg/mL, diluted 1:1 (v/v) with a buffer containing 10 mM HEPES pH 7.8, 10% (v/v) glycerol, 1 mM EDTA, 1 mM DTT and incubated 1h at 4 °C with PMAL-C8 (Anatrace, mass ratio protein:PMAL-C8 1:2.25). 30 mg BioBeads (BioRad) and TEV protease (laboratory stock, 0.6 mg/mL in 500 mM NaCl, 60% (v/v) glycerol, added at a 1:6 v/v ratio) were added, and the samples were incubated 30 min at rt followed by an overnight incubation at 4 °C. BioBeads were removed by centrifugation, and the samples were exchanged into EM amphipol buffer (10 mM HEPES pH 7.8, 85 mM NaCl, 10% glycerol, 1 mM EDTA, 1 mM DTT) using a HiTrap column (GE Healthcare). The resulting eluate was passed over 1 mL Streptactin XT high capacity beads (IBA) on a gravity flow column and washed with 5 CV EM amphipol buffer to remove non-cleaved SPC. The flowthrough was collected, concentrated and applied to size exclusion chromatography on a Superdex 200 increase 10/300 column (GE Healthcare) equilibrated with size exclusion buffer (10 mM HEPES pH 7.8, 85 mM NaCl, 1 mM EDTA, 1 mM DTT) (**Fig. S17**). Peak fractions were combined and concentrated again if necessary.

#### Activity assay

Activity assays are based on a commonly used *in vitro* cleavage protocol (52, 53). ^35^S-labeled pre-β-lactamase was generated by *in-vitro* translation. Per 5 μL reaction, 25 ng *E*.*coli* pre-ß-lactamase mRNA (Promega) were incubated with 2.5 µL rabbit reticulocyte lysate (Green Hectares), 0.5 mM DTT, and 1 µL Express ^35^S protein labelling mix (Perkin Elmer) in the presence of 0.1% (w/v) digitonin. *In vitro-*translation was allowed to proceed for 60 min at 30 °C. 2 μL of *in vitro* translated pre-β-lactamase were then incubated with 2 µL SPC (1.0 mg/mL) at 25 °C for 90 min. The samples were denatured for 15 min at 70 °C in reducing SDS-sample buffer and resolved on a 15% Tris-glycine SDS-PAGE. Gels were dried and exposed to Kodak MR films (Kodak) over night. ^35^S signal collected on the phosphor screens was scanned using a Typhoon FLA-7000 scanner (GE Healthcare). Due to the poor accessibility of the substrate signal peptide *in vitro*, partial cleavage of the precursor protein is expected (52).

#### Differential scanning fluorimetry

Melting profiles were acquired using a Prometheus NT.48 (NanoTemper). Experiments were performed using standard capillaries and a sample volume of 12 μl per capillary. SPC samples at 1.0 mg/mL were heated from 20 °C to 90 °C with 1 °C/min. UV absorbance at 350 and 330 nm were recorded at 10% excitation power. To determine the melting onset (T_on_) and melting point (T_m_), the shift in native tryptophan fluorescence was monitored by plotting changes in the emission at 350 and 330 nm. T_on_ and T_m_ were determined automatically using PR.ThermControl (NanoTemper). All experiments were performed in duplicates.

#### Mass Spectrometry

##### Shotgun mass spectrometry

###### Sample Preparation

Prior to lysis, cells were stored at -80 °C. Following mild centrifugation, the cell pellets were resuspended at a concentration of 1e6 cells per 100 µL of ice-cold lysis buffer containing 8 M urea, 50 mM Tris (pH 8), and EDTA-free mini protease inhibitor cocktail cOmplete (Sigma-Aldrich). The resuspended cell mixture was vortexed gently for 10 min and sonicated for five cycles of 30 s with a Bioraptor Plus (Diagenode SA) at 4°C. The final protein concentration was measured using a Pierce BCA protein assay kit (Thermo Fisher Scientific). An aliquot of ∼50 µg cell lysate was reduced with freshly dissolved 10 mM DTT for 30 min at 37 °C and alkylated with freshly dissolved iodoacetamide (IAA; Sigma-Aldrich) for 30 min at 37 °C in the dark. LysC (Wako Chemicals) was added at a 1:50 (w/w) enzyme-to-protein ratio and the mixture was incubated for 3 h at 37 °C. Next, the mixture was diluted to 2 M final urea concentration with 50 mM ammonium bicarbonate. Trypsin (Sigma-Aldrich) was added in 1:20 (w/w) enzyme-to-protein ratio. Digestion was performed overnight at 37 °C, followed by quenching with 0.1% trifluoroacetic acid (TFA). The digest was desalted using a Sep-Pac C_18_ 1cc vacuum cartridge (Waters). The cartridge was washed twice with 100% acetonitrile (ACN) and twice with 0.1 M acetic acid prior to sample loading. Elution was done with 80% (v/v) acetonitrile (ACN) and 0.1 M acetic acid in Milli-Q water. The desalted peptides were lyophilized by vacuum centrifugation to near-complete dryness. The final peptide mixture was resuspended in 2% (v/v) formic acid prior to LC-MS/MS data acquisition.

###### LC-MS/MS data acquisition

All data were acquired using a Thermo Scientific Ultimate 3000 RSLCnano system coupled on-line to an Orbitrap HF-X mass spectrometer (54, 55) (Thermo Scientific). Peptides were first trapped on the trapping cartridge (PepMap100 C_18_, 5 μm, 5 mm × 300 μm; Thermo Scientific) prior to separation on an analytical column (Poroshell EC-C_18_, 2.7 μm, 50 cm × 75 μm; packed in-house), heated to 40 °C. Trapping was performed for 1 min in solvent A (0.1% v/v formic acid in water), and the gradient was as follows: 9-13% solvent B (0.1% v/v formic acid in 80% v/v ACN) over 1 min, 13-44% solvent B over 95 min, 44-99% solvent B over 3 min, and finally 99% B for 4 min (flow was set to 300 nL/min). Mass spectrometry data was collected in a data-dependent fashion with survey scans from *m/z* 300 to 1500 Th (resolution of 60,000 at *m/z*=200 Th), and up to 15 of the top precursors selected and fragmented using higher-energy collisional dissociation (HCD) with a normalized collision energy of value of 27%. The MS2 spectra were recorded at a resolution of 15,000 (at *m/z*=200 Th). The AGC targets for both MS and MS2 scans were set to standard within a maximum injection time of 50 and 35 ms, respectively.

###### Data Analysis

Raw data were processed using the MaxQuant computational platform (56) with standard settings applied. In short, the extracted peak lists were searched against the reviewed Human UniProtKB database (date 15-07-2020; 20353 entries), with an allowed precursor mass deviation of 4.5 ppm and an allowed fragment mass deviation of 20 ppm. MaxQuant by default enables individual peptide mass tolerances, which was used in the search. Cysteine carbamidomethylation was set as static modification, and methionine oxidation and N-terminal acetylation as variable modifications. The iBAQ algorithm was used for calculation of approximate abundances for the identified proteins (57), which normalizes the summed peptide intensities by the number of theoretically observable peptides of the protein.

##### Native mass spectrometry

###### Sample Preparation

Samples were stored at -80 °C in either digitonin or amphipol (PMAL-C8)-containing buffer prior to native MS analysis. Approximately 10-20 µg of the membrane protein complex was concentrated and buffer-exchanged into 150 mM aqueous ammonium acetate (pH 7.5) with 0.01% (w/v) DDM, by using gel filtration with P-6 Bio-Spin columns (BioRad). The resulting protein concentration was estimated to be ∼1-2 µM before native MS analysis.

###### Data acquisition

Samples containing the membrane protein complex were directly infused into a Q Exactive Ultra High Mass Range Orbitrap instrument (QE-UHMR) (Thermo Fisher Scientific, Bremen) by using in-house prepared gold-coated borosilicate capillaries. Mass spectrometer parameters were used as follows: capillary voltage at 1.2 kV, positive ion mode, source temperature at 250 °C, S-lens RF level at 60, injection time at 50 ms, noise level parameter at 3.64. To release the membrane proteins from the detergent micelles, in-source trapping was used with a desolvation voltage of -200 V without additional collisional activation. Automatic gain control (AGC) mode was set to fixed. Resolution at 8,750 (at *m/z* = 200 Th), which corresponds to a 32 ms transient. Ion transfer optics and voltage gradients throughout the instrument were manually tuned to achieve optimal transmission of the membrane protein complex. Nitrogen was used in the higher-energy collisional dissociation (HCD) cell with trapping gas pressure setting set to 3, which corresponds to ∼2.2e-10 mBar ultra-high vacuum (UHV). The instrument was calibrated in the *m/z* range of interest using an aqueous cesium iodide solution. Acquisition of spectra was performed by averaging 1000 µscans in the time domain and subsequently recording 10 scans (2 µscans each). Peaks corresponding to the protein complex of interest were isolated with a 10 Th isolation window and probed for fragmentation using elevated HCD voltages, HCD direct eV setting of 100-150 V.

###### Data Analysis

Raw native spectra were deconvoluted with UniDec (58) to obtain zero-charged masses. For annotation, masses of the ejected subunits obtained upon activation of the membrane protein complexes were matched to the subunits identified in top-down LC-MS/MS experiment. For the reconstruction of the native MS spectrum from top-down MS data, distinct proteoforms of SPC22/23 and SEC11A or SEC11C proteins were randomly combined to obtain the masses of dimers and the products of corresponding abundances were used as abundances of the dimers. Final reconstructed native spectra were overlaid with respective native spectra obtained for the catalytic dimers of SPC-A and SPC-C complexes.

##### Intact mass and top-down mass spectrometry

###### Sample Preparation

Samples stored in digitonin or amphipol-containing buffer were diluted to a final concentration of 0.2 µg/µL. Approximately 1 µg of sample was injected for a single top-down LC-MS(/MS) experiment.

###### LC-MS(/MS) data acquisition

Chromatographic separation was achieved by using a Thermo Scientific Vanquish Flex UHPLC instrument coupled on-line, via a 1 mm × 150 mm MAbPac reversed-phase analytical column, to an Orbitrap Fusion Lumos Tribrid mass spectrometer (Thermo Fisher). The column compartment and the column preheater were heated to 80 °C during analysis. Membrane proteins were separated over a 22 min LC-MS/MS run at a flow rate of 150 µL/min. Gradient elution was performed using mobile phases A (Milli-Q water/0.1% formic acid) and B (acetonitrile/0.1% formic acid) with 30 to 57% B ramp-up in 14 min. LC-MS(/MS) data were collected with the mass spectrometer set to Intact Protein / Low Pressure. Two acquisition approaches were used with complementary full MS resolutions of either 7,500 or 120,000 (both at *m/z* = 200 Th). At 7,500 ions with masses above ∼30 kDa can be detected and at 120,000 ions with masses below ∼30 kDa can be resolved with accurate masses. Full MS scans were acquired for a mass range of *m/z* 500-3,000 Th with the AGC target set to 250% with a maximum injection time of 50 ms for the resolution of 7,500 and 250 ms for the resolution of 120,000. A total of 2 µscans were averaged and recorded for the 7,500 resolution scans and 5 µscans for the 120,000 resolution scans. All MS/MS scans were acquired with a resolution of 120,000, a maximum injection time of 250 ms, an AGC target of 10,000% and 5 µscans for the most or the first 2 most intense proteoform(s) in each cycle for medium and high resolution, respectively. The ions of interest were mass-selected by quadrupole isolation in a *m/z* = 4 Th window and collected to an AGC Target of 5e6 ions prior to electron transfer dissociation (ETD). The ETD reaction time was set to 16 ms with a maximum injection time of 200 ms and the AGC target of 1e6 for the ETD reagent. For data-dependent MS/MS acquisition strategy, the intensity threshold was set to 2e5 of minimum precursor intensity. MS/MS scans were recorded in the range of *m/*z = 350-5000 Th using high mass range quadrupole isolation.

###### Data Analysis

Full MS spectra were deconvoluted with either Xtract or ReSpect (Thermo Fisher Scientific) for isotopically-resolved or unresolved data, respectively. Automated proteoform searches against a custom sequence database were performed in Thermo Proteome Discoverer (version 2.4.0.305) extended with the ProSightPD 3.0 nodes (59). Parameters were set as follows. ReSpect: precursor *m/z* tolerance – 0.2 Th; relative abundance threshold – 0%; precursor mass range – 3-100 kDa; precursor mass tolerance – 30 ppm; charge range – 3-100. Xtract: signal/noise threshold – 3; *m/z* range – 500-3,000 Th. Initially, a large precursor tolerance window of 5 kDa was used to allow for detection of unknown PTMs and sequence processing followed by cycles of database filtering and sequence adjustment to determine a final set of isoforms/proteoforms. For the final database search, ProSightPD parameters were: precursor mass tolerance – 500 Da; fragment mass tolerance – 20 ppm. To verify unreported isoforms/proteoforms, in-house R and C# scripts were used to group replicate fragmentation scans for a precursor of interest followed by automated fragment annotation and manual spectrum inspection. A similar approach was used to characterize unidentified abundant precursors (60). Representation of proteoforms per protein was achieved by summing full MS scans per protein elution peak and converting spectra to zero-charged mass profiles in UniDec (58).

##### Cross-linking mass spectrometry

###### Sample Preparation

Proteins were incubated with the cross-linking reagent PhoX (24) for 45 min at room temperature (buffer conditions specified below, **Fig. S17**). The cross-linking reaction was quenched by addition of Tris·HCl (100 mM, pH 7.5) to a final concentration of 10 mM. Cross-linked proteins were further purified from aggregation products by size exclusion chromatography on a Superose 6 increase column (GE Healthcare) equilibrated with size exclusion buffer (10 mM HEPES pH 7.8, 85 mM NaCl, 1 mM EDTA, 1 mM DTT, 0.09% (w/v) digitonin). The concentration of peak fractions ranged between 0.2 mg/mL (SPC-A) to 0.4 mg/mL (SPC-C). Crosslinked proteins were denatured by addition of urea (8 M in 100 mM Tris) and reduced by addition of DTT (final concentration of 2 mM) for 30 min at 37 °C, followed by alkylation with IAA (final concentration of 4 mM) for 30 min at 37 °C. Afterwards the sample was digested by incubation with a combination of LysC (1:75 enzyme to protein) and Trypsin (1:50 enzyme to protein) for 10 h at 37 °C, after which formic acid (final concentration 1%) was added to quench the digestion. Finally, peptides were desalted by Sep-Pak C_18_ prior to Fe-IMAC enrichment.

Cross-linked peptides were enriched with Fe(III)-NTA cartridges (Agilent Technologies) using the AssayMAP Bravo Platform (Agilent Technologies) in an automated fashion. Cartridges were primed at a flow rate of 100 μL/min with 250 μL of priming buffer (0.1% TFA, 99.9% ACN) and equilibrated at a flow rate of 50 μL/min with 250 μL of loading buffer (0.1% TFA, 80% ACN). The flow-through was collected into a separate plate. Dried samples were dissolved in 200 μL of loading buffer and loaded at a flow rate of 5 μL/min onto the cartridge. Cartridges were washed with 250 μL of loading buffer at a flow rate of 20 μL/min and cross-linked peptides were eluted with 35 μL of 10% ammonia directly into 35 μL of 10% formic acid. Samples were dried down and stored at -20 °C prior to further use. Before to LC–MS/MS analysis, the samples were resuspended in 10% formic acid.

###### LC-MS/MS data acquisition

All data were acquired using an UHPLC 1290 system (Agilent Technologies) coupled on-line to an Orbitrap Fusion (61) mass spectrometer (Thermo Scientific). Peptides were first trapped (Dr. Maisch Reprosil C_18_, 3 μm, 2 cm × 100 μm) prior to separation on an analytical column (Agilent Poroshell EC-C_18_, 2.7 μm, 50 cm × 75 μm). Trapping was performed for 10 min in solvent A (0.1% v/v formic acid in water), and the gradient for protein complexes was as follows: 0 – 10% solvent B (0.1% v/v formic acid in 80% v/v ACN) over 5 min, 10-40% solvent B over 70 min, 40-100% solvent B over 3 min, and finally 100% B for 4 min (flow was passively split to approximately 200 nL/min). The mass spectrometer was operated in a data-dependent mode. Full-scan MS spectra were collected in a mass range of *m/z* 350 – 1300 Th in the Orbitrap at a resolution of 60,000 after accumulation to an AGC target value of 1e6 with a maximum injection time of 50 ms. In-source fragmentation was activated and set to 15 eV. The cycle time for the acquisition of MS/MS fragmentation scans was set to 3 s. Charge states included for MS/MS fragmentation were set to 3-8 respectively. Dynamic exclusion properties were set to n = 1 and to an exclusion duration of 20 s. HCD fragmentation (MS/MS) was performed in stepped collision energy mode (31.5, 35, 38.5 %) in the Ion Trap and the mass spectrum acquired in the Orbitrap at a resolution of 30,000 after accumulation to an AGC target value of 1e5 with an isolation window of *m/z* = 1.4 Th and maximum injection time of 120 ms.

###### Data Analysis

The acquired raw data were processed using Proteome Discoverer (version 2.4.0.388) with the XlinkX/PD nodes integrated (24, 62). The linear peptides search was performed using the standard Mascot node as the search engine with the full Human database from UniProtKB (20,230 entries, downloaded from UniProtKB downloaded at 2018_01). Cysteine carbamido-methylation was set as fixed modification. Methionine oxidation and protein N-term acetylation was set as dynamic modification. For the search of mono-links, water-quenched (C_8_H_5_O_6_P) and Tris-quenched (C_12_H_14_O_8_PN) were set as dynamic modifications. Trypsin/P was specified as the cleavage enzyme with a minimal peptide length of six and up to two miss cleavages were allowed. Filtering at 1% false discovery rate (FDR) at the peptide level was applied through the Percolator node. For crosslinked peptides, a database search was performed against a FASTA containing the proteins under investigation supplemented with a common contaminants list of 200 proteins using XlinkX/PD nodes for cross-link analysis. Cysteine carbamidomethylation was set as fixed modification and methionine oxidation and protein N-term acetylation were set as dynamic modifications. Trypsin/P was specified as enzyme and up to two missed cleavages were allowed. Furthermore, identifications were only accepted with a minimal score of 40 and a minimal delta score of 4. Otherwise, standard settings were applied. Filtering at 1% false discovery rate (FDR) at peptide level was applied through the XlinkX Validator node with setting simple.

##### Single Particle analysis

###### Sample preparation

Size exclusion chromatography (SEC) peak fractions were either used directly for vitrification (at 0.5 mg/mL) or concentrated to 4 mg/mL and vitrified in the presence of 1.5 mM fluorinated fos-choline (FFosC, Anatrace). 3 µL samples were applied to freshly glow-discharged Cu 200 Holey Carbon R1.2/1.3 grids (Quantifoil, for samples without FFosC), or Cu 200 Holey Carbon R2/1 grids (Quantifoil, for samples containing 1.5 mM FFosC). In both cases, grids were flash-frozen using a Vitrobot Mark IV (Thermo Fischer Scientific) with 595 blotting paper (Ted Pella) at 4 °C, 100% humidity, and either a blot force of 0 for 4 s or a blot force of -2 for 3 s and a liquid ethane/propane mixture.

###### Data collection

Data were collected on a 200 kV Talos Arctica microscope (Thermo Fischer Scientific) equipped with a post-column energy filter (slit width 20 eV) and a K2 summit direct electron detector (Gatan). EPU (Thermo Fischer Scientific) was used for automated data collection in counted mode. Movies were acquired in 45-50 frames at an effective pixel size of 0.81 Å/px, with a dose rate of ∼4 e^-^/px/s (measured in an empty hole without ice), and a total dose of 60 e^-^/Å^2^. Defocus values ranged between 0.5 and 4 μm. Data quality was monitored in real time using Warp (63).

###### Image processing

All four datasets collected for SPC-A and SPC-C in PMAL-C8 were processed analogously (for detailed dataset statistics, refer to **Table S1** and **Fig. S3-S4**). Collected movie stacks were manually inspected and imported into Relion 3.1 (64). Motion correction was performed with MotionCor2 (65), and CTFFIND4 (66) was used for CTF estimation using exhaustive search in Relion 3.1. Movies with an estimated resolution worse than 10 Å were discarded. For particle picking, the motion-corrected micrographs were denoised using a trained model in SPHIRE-JANNI (67). Particle picking was performed with SPHIRE-crYOLO 1.5.5 (68), using models trained separately on subsets with similar defocus (0-1 μm, 1-2 μm, 2-3 μm, and 3-4 μm) to ensure maximal particle recovery across the whole defocus range. Particles were extracted in Relion 3.1 at 3-fold binning and subjected to 3 rounds of 2D classification, during which information until the first CTF zero was ignored (**Fig. S3**). Once 2D classification converged, the respective datasets with and without FFosC were combined and subjected to 2 additional rounds of 2D classification. Unbinned images of the remaining particles were used to generate an initial 3D model, followed by 3D refinement, and re-extraction with updated coordinates. Since the micelle occupies about 50% of the particle volume, it was expected to have detrimental effects on particle alignment. Therefore. the refined map was used as a reference for a 3D classification without particle alignment using a mask enclosing only the protein portion. The respective best classes were subjected to CTF correction, Bayesian polishing, and 3D refinement in Relion 3.1. Further attempts to perform CTF corrections and subtract the micelle did not improve the reconstructions. A post-processing step in Relion was employed to partially mask the micelle, correct with the detector MTF, and apply a sharpening B-factor of -180 Å^2^.

###### Model Building & Refinement

All models were generated by subjecting the full-length proteins to structure prediction by trRosetta (21) (**Fig. S5**). The precise pixel spacing was determined by map correlation of the SEC11 luminal domain in UCSF Chimera (69). Models for SPC12 and SPC25 were docked as complete rigid bodies, whereas models of SEC11A and SEC11C were separated into two rigid body groups: (i) the N-terminal part containing the cytosolic portion and the single TM helix, and (ii) the luminal domain including the C-terminus. Both groups were docked individually in UCSF Chimera. Similarly, SPC22/23 was divided into three rigid body groups: (i) the N-terminal TM helix, (ii) a short strand with poor density fit that connects the TM helix and the rest of the protein, and (iii) the luminal domain. An additional model containing only the luminal portion was calculated and used to mitigate influences of the TM helix on the fold of the luminal portion. All domains except SPC12 could be docked unambiguously based on their soluble parts (**Fig. S5**). SPC12 was fitted based on the different lengths of its two TM segments and the presence of a hydrophilic stretch that is unlikely to be exposed to the membrane interface. Existing information in the literature, top-down, native, and XL-MS data, the presence of known anchor points (glycosylation site at Asp141 of SPC22/23, visible tryptophan side chains), secondary structure predictions, disorder predictions, atomistic molecular dynamics simulations of the protein in POPC using a CHARMM force field (see below), differences in the top-5 trRosetta models, and overall biochemical properties of the resulting model were used to evaluate the docking and domain architecture (**Fig. S6**).

Unresolved parts of the assembled SPC were trimmed manually in *Coot* version 0.9 (70) (**Fig. S5**). Alternating rounds of manual adjustment in *Coot* and PHENIX real space refine (71) were applied to yield the final models. In *Coot*, all-atom restraints and tight geometry restraints were used at all times. Only loops with poor density fit and, if necessary, side chains with low-probability rotamers were adjusted. In PHENIX, morphing and simulated annealing were applied to enable fitting of loops. Map-model FSCs were calculated using PHENIX Mtriage. All figures were prepared using UCSF ChimeraX (72) or PyMol (Schrödinger).

##### Surface potential calculation

Surface potentials were calculated in vacuum using the APBS plugin in PyMol (Schrödinger), based on pKa calculations using the built-in pdb2pqr module.

##### Secondary structure and disorder prediction

Secondary structure elements were predicted using JPred4 (73). Disordered regions were predicted using the HHblits method with NetSurfP-2.0 (74)

##### Phylogenetic analysis and residue conservation

Representative, curated protein sequences of SPC subunits from all major branches of life were manually extracted from UniProtKb. Sequence alignments and phylogenetic trees were computed using MUSCLE in Mega-X (75). Residue conservation was calculated using the Consurf server (76) searching against the cleaned UniProtKB database using the HMMER algorithm.

##### Molecular dynamics simulations

###### Coarse-grained models

The most recent development version of the Martini 3 Coarse-Grained (CG) force field (77) was used to perform all CG molecular dynamics (MD) simulations. The CG protein model was generated with the new version of the program Martinize (78, 79). The refined full-length SPC-C model generated by trRosetta and fitted into the observed density was used as a reference to define bonded parameters dependent of the secondary structure. The glycan chain attached to Asp141 of SPC22/23 was not included in the model. Elastic networks were applied to each monomer of the SPC-C complex, with a distance cut-off of 0.85 nm using a force constant of 1300 kJ mol^−1^ nm^−2^. Unresolved parts of SPC-C were kept free, without any elastic bonds. Additional harmonic bonds were added between the protein monomers, to further increase stability of the SPC-C complex. With a cut-off of 0.7 nm, these extra harmonic potentials mimic hydrogen bonds between the backbone beads of different monomers. Lipids models were inspired on the previous Martini 2 force-field (80, 81), but now following the rules of Martini 3 and with adaptations in the bonded parameters inspired by the “extensible model” of Carpenter *et al*. 2018 (82).

###### System setup and settings of the coarse-grained MD simulations

SPC-C was embedded in different membrane environments including an endoplasmic reticulum membrane model, as described in **Table S3**. All systems were solvated using a Martini water model solution with 0.15 M concentration of NaCl, mimicking physiological conditions. All simulation boxes were built using the INSANE program with dimensions of 18 × 18 × 15 nm^3^ (81). The principal axis of the SPC-C complex was set to be parallel to the normal of the lipid bilayers. Firstly, the system was minimized for 2000 steps with the steepest descent method, followed by an equilibration stage performed for 500 ps with 10 fs as time step. After minimization and equilibration, the production run was performed for 20 µs, using a time step of 20 fs. Settings for the CG simulations followed the “new” Martini set of simulation parameters (83). The temperature of the systems was kept at 310 K with the velocity rescaling thermostat (84). For the pressure, we used semi-isotropic coupling at 1 bar using the Parrinello-Rahman barostat (85). Additional MD simulations of pure lipid bilayers were performed to be compared with SPC-C embedded in the same environments. All simulations were performed with GROMACS (version 2020) (86).

###### System setup and settings of atomistic MD simulations

The CHARMM force field for proteins (87) and lipids (88) was used to perform the atomistic MD simulations. Water was modelled explicitly using the modified CHARMM TIP3P model (89, 90). The temperature was coupled to a heat bath at 310 K, using the velocity rescale thermostat (84). The pressure was kept at 1.0 bar, using a Parrinello−Rahman barostat (85) with a compressibility of 4.5 × 10^−5^ bar^−1^and coupling time of 4.0 ps. Particle Mesh Ewald (91, 92) was used to compute the electrostatic interactions, with a real-space cut-off of 1.2 nm. Van der Waals interactions were switched to zero between 1.2 and 1.4 nm. Neighbor lists were updated every 10 steps. Bonds involving hydrogens were constrained using the LINCS algorithm (93). The integration time step used was 2 fs and the overall center of mass motion was removed every 10 steps. Simulation box and topologies were initially built with CHARMM-GUI (94). SPC-C was embedded in a POPC bilayer and solvated with water in a simulation box of 18 × 18 × 15 nm^3^. NaCl was added to bring the system to neutral charge and an ionic strength of 0.15 M. Minimization and equilibrations was based on the CHARMM-GUI protocol, followed by a 150 ns production run with time step of 2 fs. As the relaxation of the bilayer (which includes the thinning in the TM window) demanded longer simulations, an additional initial configuration of SPC-C in POPC was built from the final configuration of the 20 µs CG MD simulations. The CG configuration was backmapped to the atomistic resolution using the backward program (95), which was followed by another 150 ns production run. All atomistic simulations were performed with GROMACS (version 2020) (86).

###### Analysis of the trajectories

The thickness (*d*_*i*_) of the membranes were estimate based on the average distance of the phosphate bead (in CG simulations) or phosphorus atom (in atomistic simulations) of the lipids using the *gmx density tool* of GROMACS (86). The percentage of thinning (%*T*) was estimated based on the difference of thickness in the bulk membrane (*d*_*membrane*_) and the lipids inside the TM window (*dTM*_*window*_) of the SPC-C complex, according to equation (1):

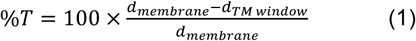

Lipid enrichment (%*E*_*i*_) near the TM window was computed according to equation (2):

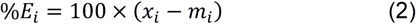

where *x*_*i*_ is molar fraction of lipids near the TM window and (*m*_*i*_) is the molar fraction of the lipids in relation of the whole membrane. *m*_*i*_ was simply defined by the composition of the membrane. On the other hand, *x*_*i*_ was defined as the number of contacts of the lipid with the TM window (*c*_*i*−*TM window*_) in relation of the total number of contacts with all the lipids, according to equation (3):

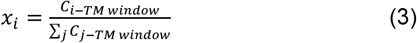

*c*_*i*− *TM window*_ was computed using the *gmx mindist tool* of GROMACS (86). A distance cut-off of 1.1 nm was used between the PO4 and ROH beads (from phospholipids and cholesterol, respectively), and the BB beads of the following residues around the transmembrane helices of SPC-C complex: Chain A - Tyr31, Tyr32, Gln36, Tyr185, Val186 and Leu,187; Chain B – Ala8, Asn9 and Ser10; Chain C – Thr129, Ile130 and Tyr131; Chain D – Glu88, Gln89 and Met90.

Root mean square fluctuation (RMSF) calculation was performed using a Fortran program, based on the MDLovoFit code (96). The alignment protocol of the Cα atoms starts with a standard rigid-body alignment of the structures using all Cα atoms as reference, followed by the calculation of the RMSF per residue for atoms. After the identification of the residues with RMSF lower than 2 Å, a new rigid body alignment uses only this subgroup of residues that represents the most rigid parts of SPC-C. This protocol is repeated until the RMSD and the residues used in the rigid subgroup are converged. As structure reference for the trajectory alignment, we used the SPC-C average structure from the trajectory.

Visual inspection and figure rendering of the trajectory snapshots were performed with VMD (97).

## Acknowledgments

Research has been supported by the ERC Consolidator Grant 724425 (Biogenesis and Degradation of Endoplasmic Reticulum Proteins, to F.F.), the research program TA with project number 741.018.201 (to R.A.S. and F.F.), which is partly financed by the Dutch Research Council (NWO), and the ERC Advanced Grant “COMP-MICR-CROW-MEM” (to S.J.M.); additional support came through the European Union Horizon 2020 program INFRAIA project Epic-XS (Project 823839). This work benefited from access to the NKI Protein Facility, an Instruct-NL and Instruct-ERIC center. P.C.T.S., and S.J.M. acknowledge the National Computing Facilities Foundation (NCF) of the NWO for providing computing time. The authors are thankful to Dr. Robert-Jan Lebbink and Dr. Patrique Praest (UMC Utrecht) as well as Prof. Dr. Thijn Brummelkamp (NKI Amsterdam) who kindly provided the cell lines used to characterize SEC11 expression. We thank Dr. Stuart Howes (Utrecht University) for EM support, Susanne Bruekner (NKI Amsterdam) for support with nanoDSF, and Panagiotis Drougkas (Utrecht University) for help with protein expression and purification. Further, we are grateful for the infrastructure provided by the cellular protein chemistry laboratory at Utrecht University, and thank Prof. Dr Ineke Braakman, Guus van Zadelhoff, Dr. Juliette Fedry and Lena Thärichen for help with activity assays. We thank Prof. Dr. Richard Zimmerman for critically reading the manuscript.

## Author Contributions

A.M.L. and F.F. conceived the project. A.M.L. designed expression constructs. A.M.L. and M.G.M. cloned constructs and point mutants. A.M.L. expressed cells and established a protein purification protocol. A.M.L. and P.O. performed protein purification and sample purification for cryo-EM and MS. B.S., S.T., R.A.S., A.M.L., and F.F. conceived MS experiments. B.S., R.A.S. and S.T. performed MS experiments and data analysis. A.M.L. and P.O. solved the structures of SPC-C and SPC-A, including sample preparation, data collection and data processing refinement. A.M.L. and F.F. generated, refined, and analyzed atomic models. A.M.L. conceived and performed stability and activity assays. P.C.T.S., A.M.L., S.J.M., and F.F. conceived MD simulation experiments, which P.C.T.S. performed and analyzed. M.G.M. grew cells for shotgun analysis. A.M.L., B.S., S.T., P.C.T.S., S.J.M., R.A.S., and F.F. wrote the manuscript and prepared the figures with help from all co-authors.

## Notes

The authors declare no competing interests.

