## Supplementary Information for "Structure of the Human Signal Peptidase Complex Reveals the Determinants for Signal Peptide Cleavage"

Liaci *et al.* 2020

**Content:**

- Supplementary Text
- Supplementary Figures 1-17
- Supplementary Tables 1-3
- Legends for Supplementary Movies 1-3

### Supplementary Text

#### Native mass spectrometry of SPC paralogs

Upon gas-phase activation, SPC dimers, overall, exhibit comparable dissociation stabilities, whereby SPC-A exhibits a slightly more stable dimer with ~45 % of total ion intensity corresponding to the ejected subunits compared to ~65 % for SPC-C at the same dissociation conditions. Additionally, SEC11A has a more stable conformation within the dimeric complex compared to SEC11C, which is estimated based on the intensity-weighted retained charge of an ejected subunit. Here, in the case of SPC-A, SEC11A and SPC22/23 retain ~32% and ~53% of their original precursor charges, respectively, upon dissociation; in the case of SPC-C, SEC11C and SPC22/23 retain ~44% and ~36% of their original precursor charges, respectively. The higher average charge of dissociated SEC11C is indicative of a more extended conformation, *i.e.* a higher degree of unfolding (98, 99), compared to SEC11A.

### Supplementary Figures

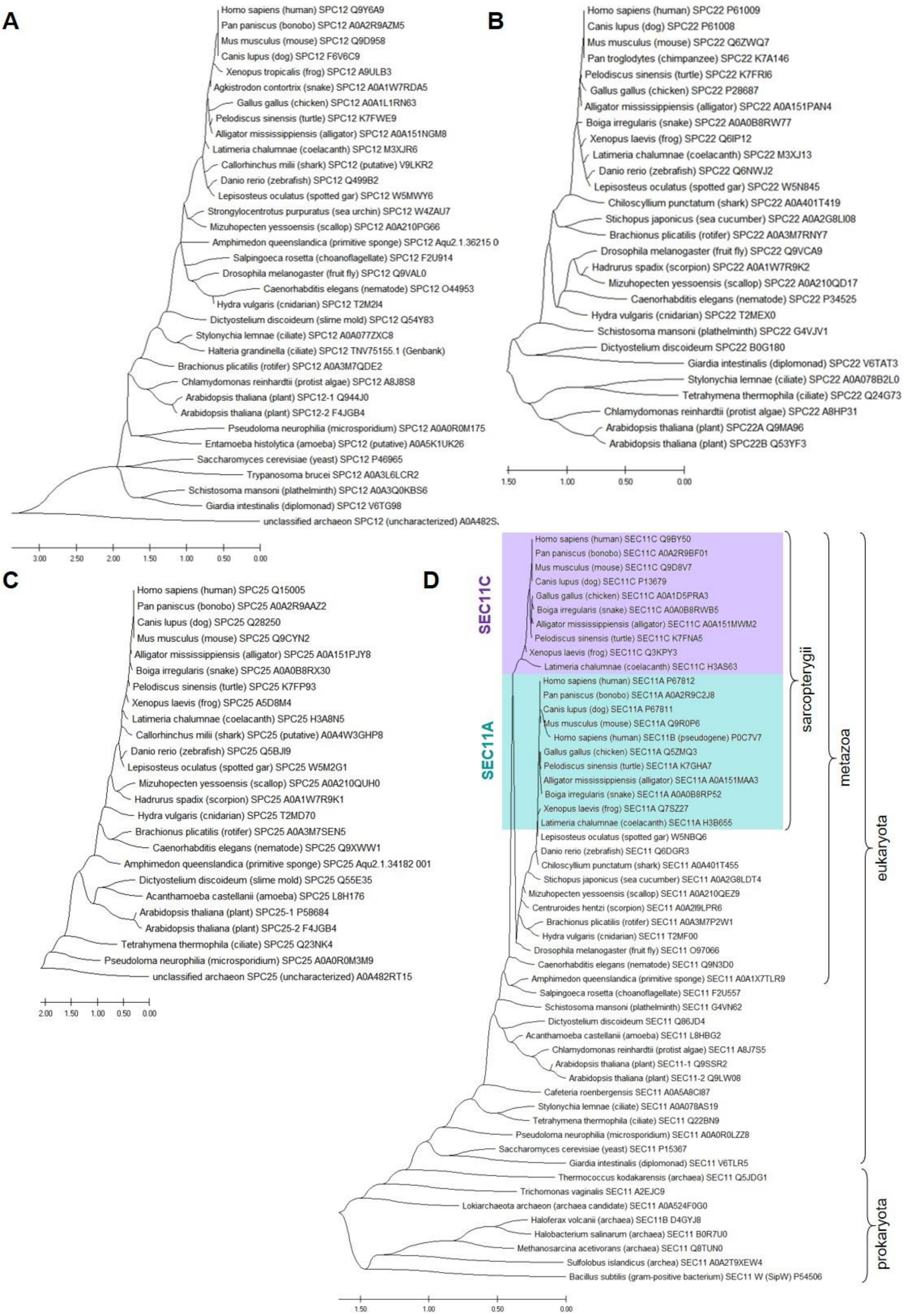


**Figure S1**. | **Phylogeny of SPC subunits.** (**A**) Phylogenetic tree of SPC12. (**B**) Phylogenetic tree of SPC22/23. (**C**) Phylogenetic tree of SPC25. (**D**) Phylogenetic tree of ER-type SPases. SEC11A sequences are highlighted in teal, SEC11C in purple. SEC11C proteins form a distinct clade. The earliest known ancestor with two SEC11 paralogs is the 'living fossil' coelacanth *Latimeria chalumnae*. The evolutionary distances (x-axis) are in the units of the number of amino acid substitutions per site.

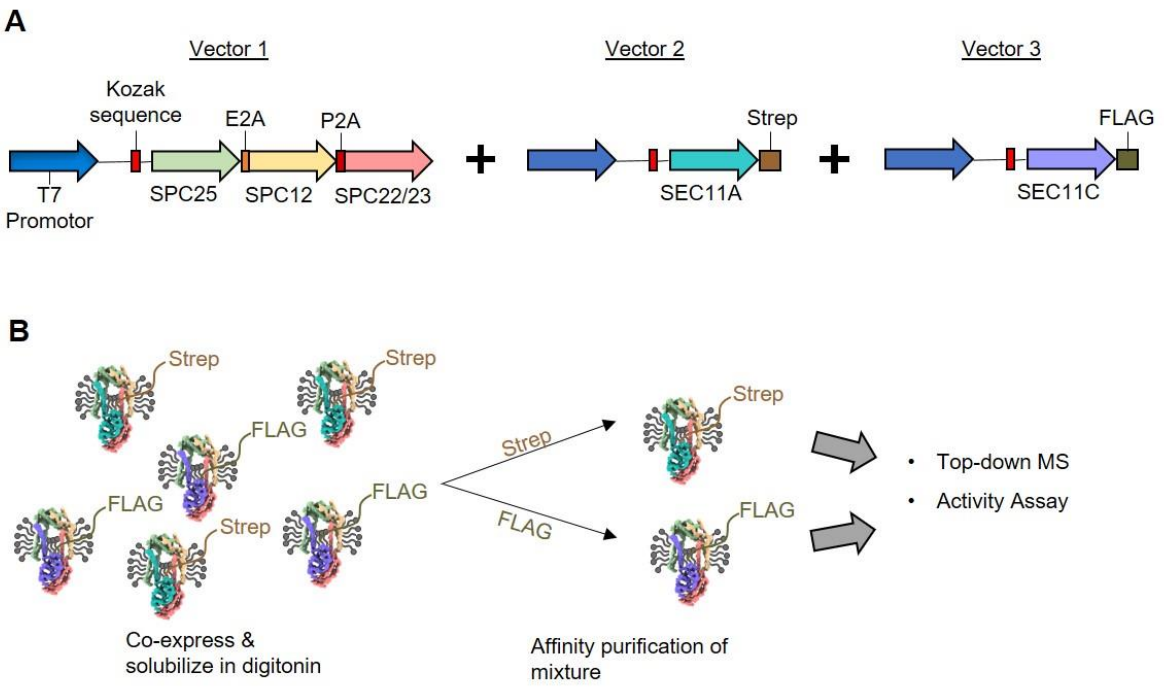


**Figure S2 | Expression constructs and purification workflow for SPC composition experiments**. (**A**) Expression constructs used for the analysis of SPC composition. (**B**) Description of the purification workflow. Accessory subunits were co-expressed in HEK 293 cells with SEC11A-Strep and SEC11C-FLAG in a vector mass ratio of 4:1:1 and solubilized in digitonin. The batch was purified with both Strep and FLAG affinity resin, and the respective eluates were analyzed for composition and *in vitro* activity.

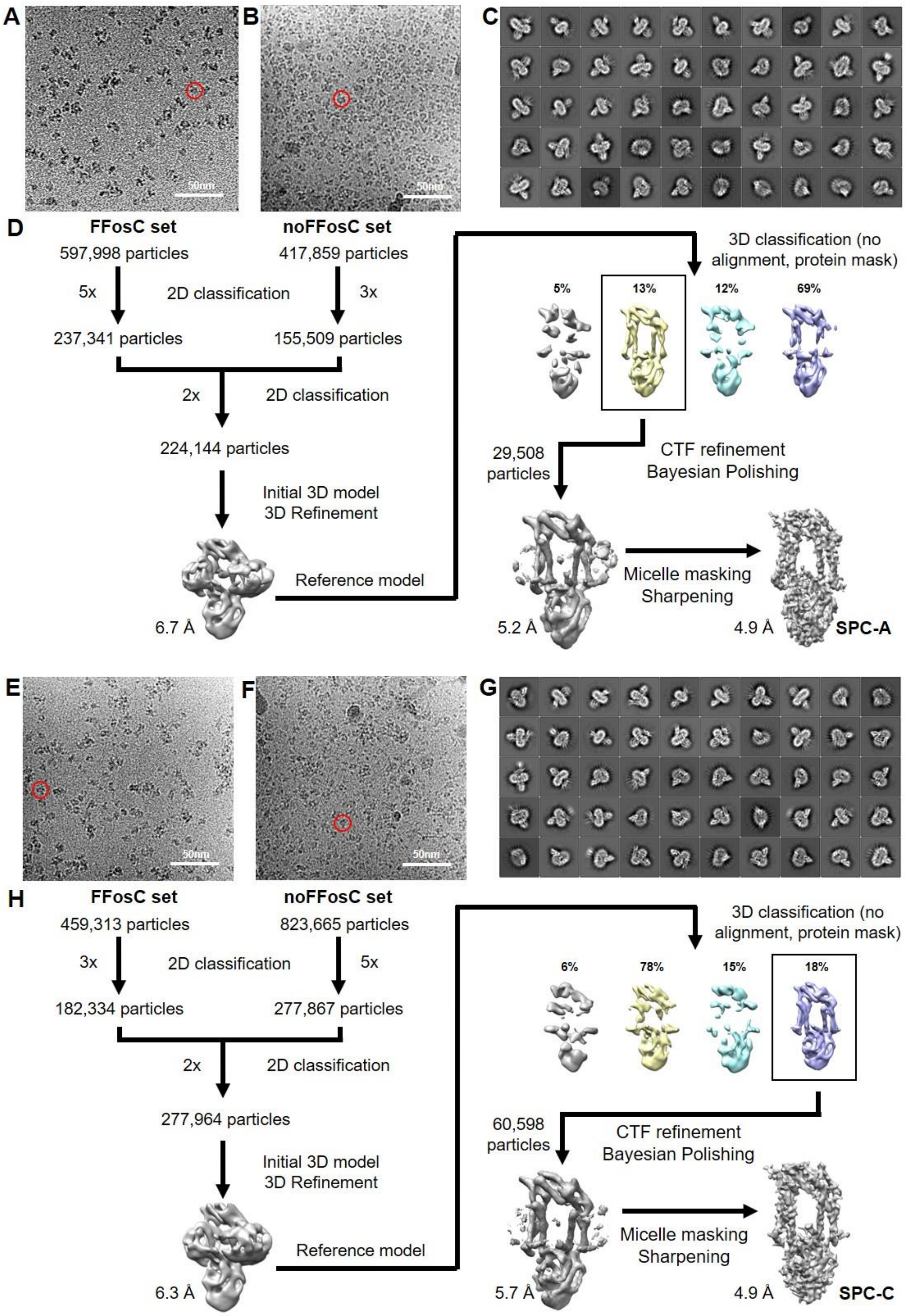


**Figure S3 | Single Particle Data Processing Workflow.** (**A**) Representative micrograph of SPC-A collected with 1.5 mM FFosC at 2.6 µm defocus. Exemplary particle highlighted by a red circle. (**B**) Representative micrograph of SPC-A collected without FFosC at 2.9 µm defocus. Protein particles are interspersed with empty amphipols. (**C**) Top 50 2D classes for the SPC-A dataset. (**D**) Processing workflow for the SPC-A dataset. (**E**) Representative micrograph of SPC-C collected with 1.5 mM FFosC at 2.6 µm defocus. (**F**) Representative micrograph of SPC-C collected without FFosC at 2.6 µm defocus. (**G**) Top 50 2D classes for the SPC-C dataset. (**H**) Processing workflow for the SPC-C dataset.

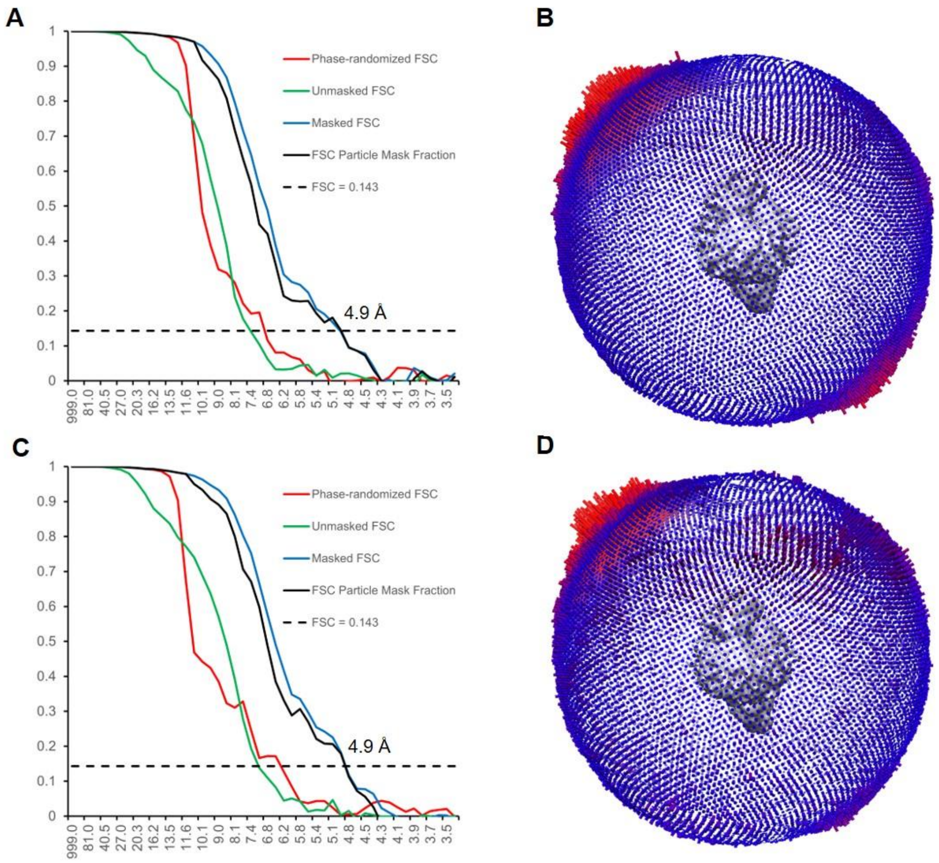


**Figure S4 | Cryo-EM map quality metrics.** (**A**) FSC curve of SPC-A as reported by Relion. (**B**) Angular distribution of particles within the SPC-A dataset (**C**) FSC curve of SPC-C as reported by Relion. (**D**) Angular distribution of particles within the SPC-C dataset.

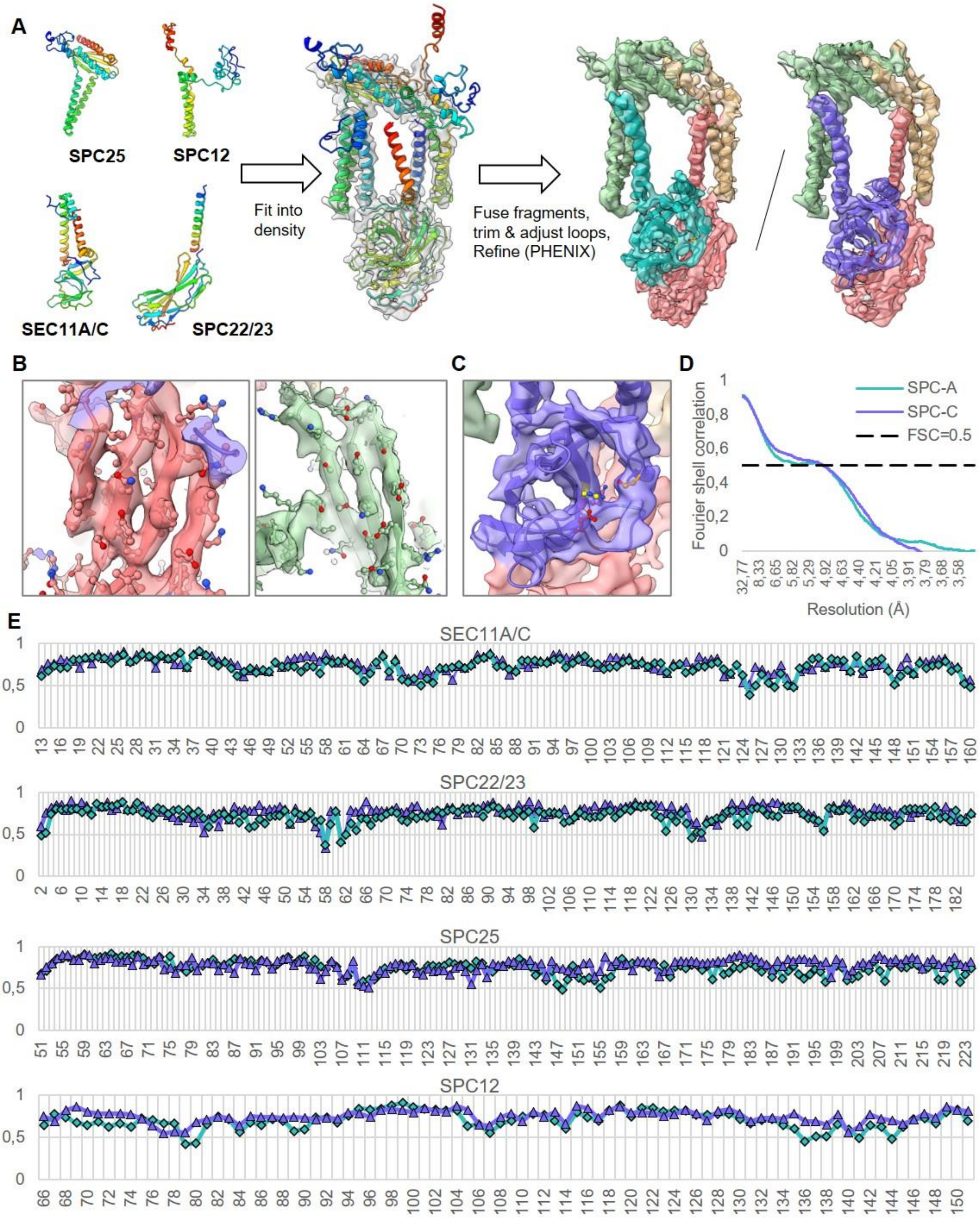


**Figure S5 | SPC model building.** (**A**) Initial structures of SPC subunits were calculated using trRosetta and full-length sequences (UniProtKb IDs Q15005, Q9Y6A9, P61009, P67812, Q9BY50). These models were fitted into the observed density maps. Flexible regions were trimmed and adjusted using all-atom refinement in Coot 0.9 with tight geometry restraints. The models were refined against the density maps using PHENIX real space refine. If necessary, rotamer outliers and regions with poor density fit were adjusted in Coot and subjected again to PHENIX real space refine. Individual subunits are colored according to their residue number (from blue to red), and the subunits of assembled SPC-A and SPC-C are colored as in Fig. 1. (**B**) Representative map features of SPC-C. Beta sheets and large, well-ordered side chains can be distinguished. (**C**) Map-to-model fit of the luminal domain of SPC-C. (**D**) Map–to-model FSC curves for SPC-A and SPC-C as determined by PHENIX Mtriage. (**E**) Per-residue real space correlation as reported by PHENIX real space refine. Teal diamonds = SPC-A subunits; purple triangles= SPC-C subunits.

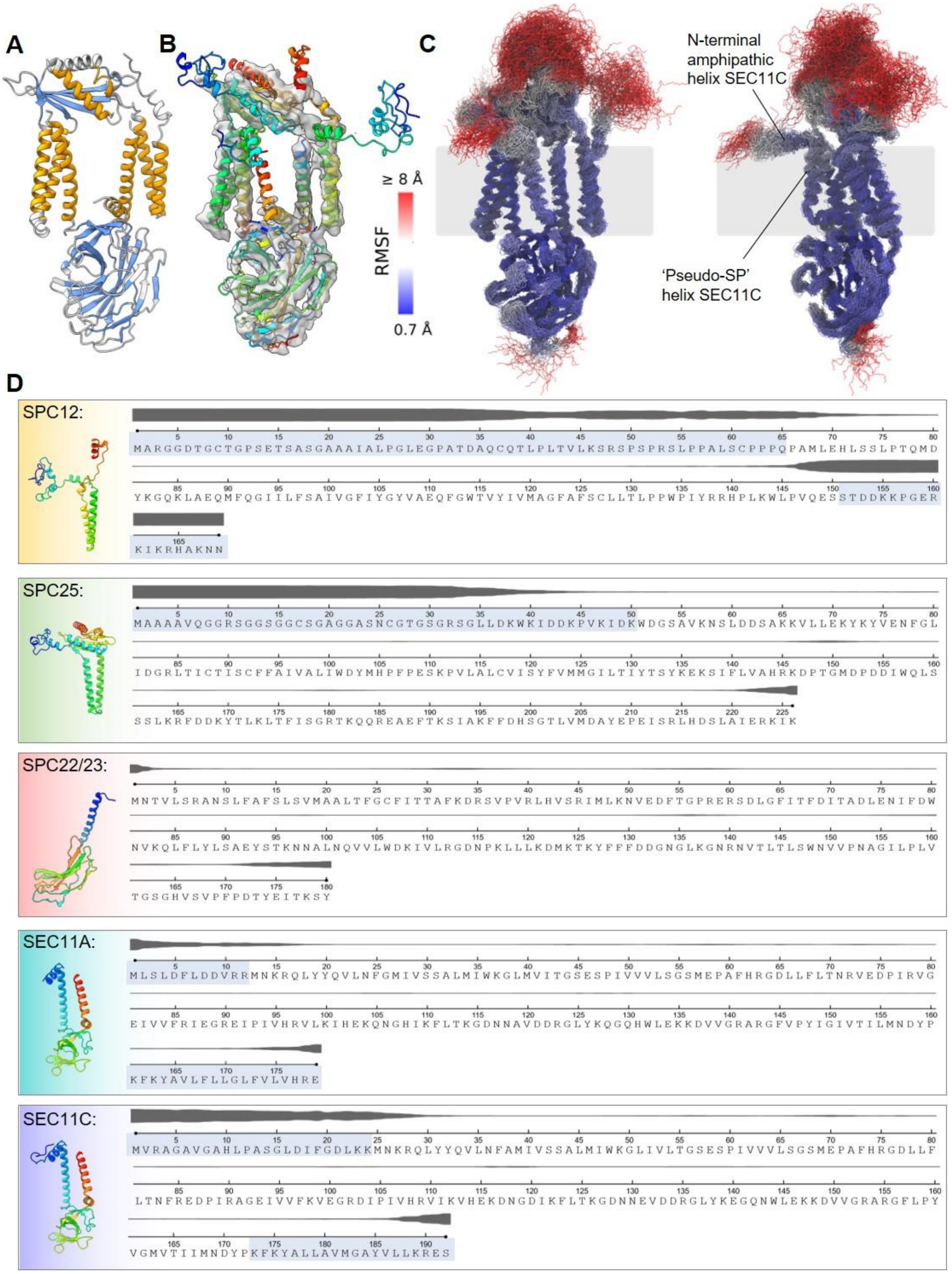


**Figure S6** | **Secondary structure and disorder predictions.** (**A**) Predicted secondary structure elements (orange = alpha helices; blue = beta strands) are mapped onto the observed SPC-C model. (**B**) Areas without ordered density are depicted. (**C**) Flexibility of the SPC-C model in atomistic MD simulations. Two views of representative aligned snapshots obtained from a 150 ns MD simulation of SPC-C embedded in POPC are shown. The color scale represents the backbone RMSF in relation to the average structure of the simulations. Termini of SEC11 and the membrane (grey transparent box) are indicated. Except for the experimentally unresolved parts, the overall organization of the SPC complex was maintained. Some small relaxation of TM helices is observed which is consistent with the different environment of experiments (amphipol micelle) and simulations (phospholipid bilayers). (**D**) Disorder predictions for each subunit are shown as grey lines. The thicker the line, the higher the disorder probability. Highlighted sequence stretches are not resolved in the atomic models. Full-length trRosetta models (colored as in Fig. S5) are shown.

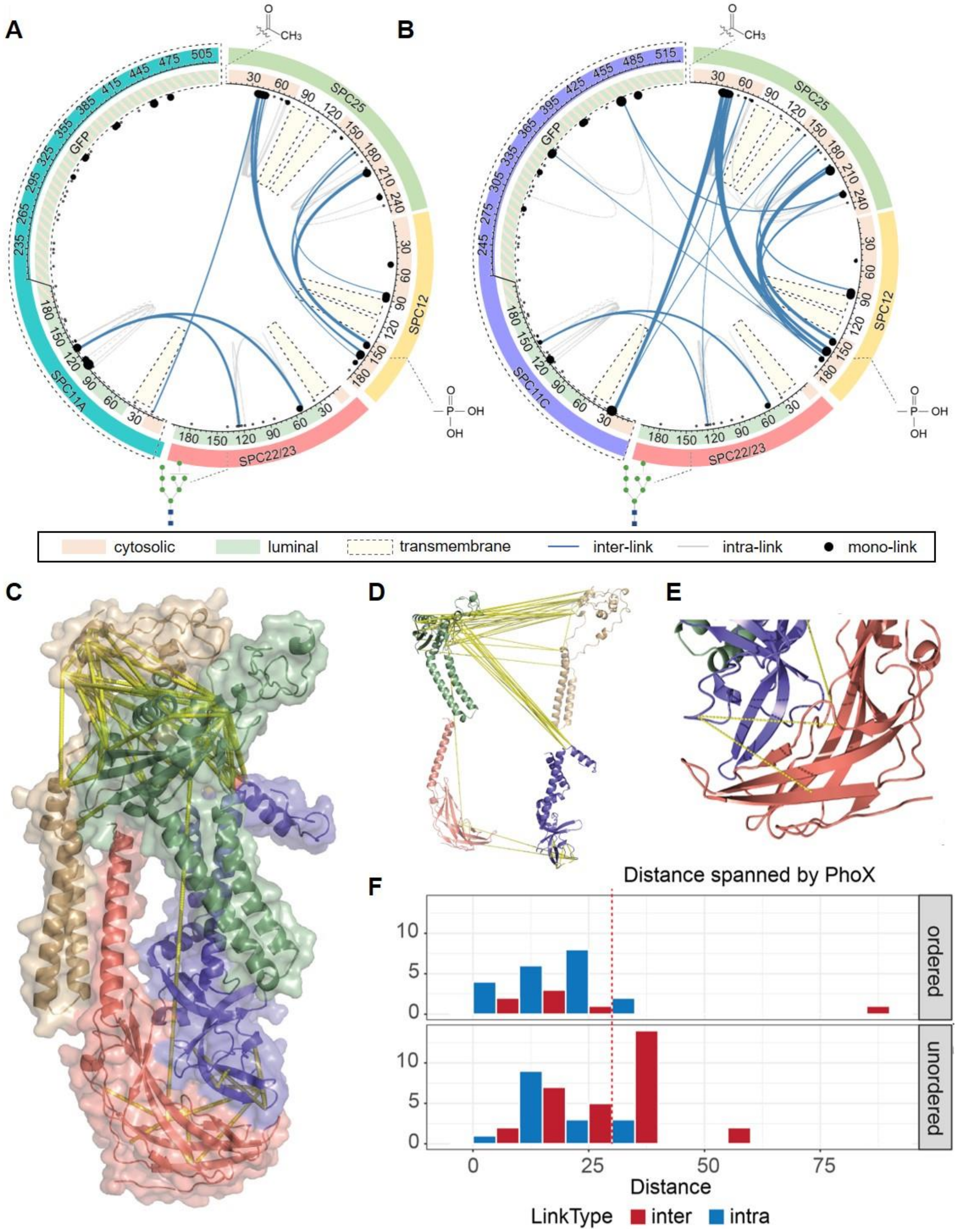


**Figure S7 | XL-MS of SPC-A and SPC-C.** Circos representation of the detected cross-links for SPC-A and SPC-C demonstrates excellent overlap in the structure. As anticipated, no cross-links are detected inside the transmembrane domains. In addition, very few cross-links are detected to the exposed GFP tag, indicating this protein region remains out of distance from the complex. (**A**) Detected cross-links for SPC-A. (**B**) Detected cross-links for SPC-C. (**C**) The identified cross-links cluster in the exposed regions of the complex with a heavy preference for the cytosol-exposed part. (**D**) The organization of the individual subunits in the complex is readily revealed by the identified cross-links. (**E**) The SPC22/23-SEC11A/C interface, as revealed by native mass spectrometry, is highly stable. Relatively few cross-links are located here due to the low flexibility and lower abundance of lysines. (**F**) The measured distances in ordered parts of the complex reveal that the intra-links are well within the distance constraint determined for the PhoX crosslinker (24) (indicated by a red dashed line). The distances for the disordered regions are an estimation based on the structural model predicted by trRosetta. These regions were not detected in the EM density and likely represent a high degree of error.

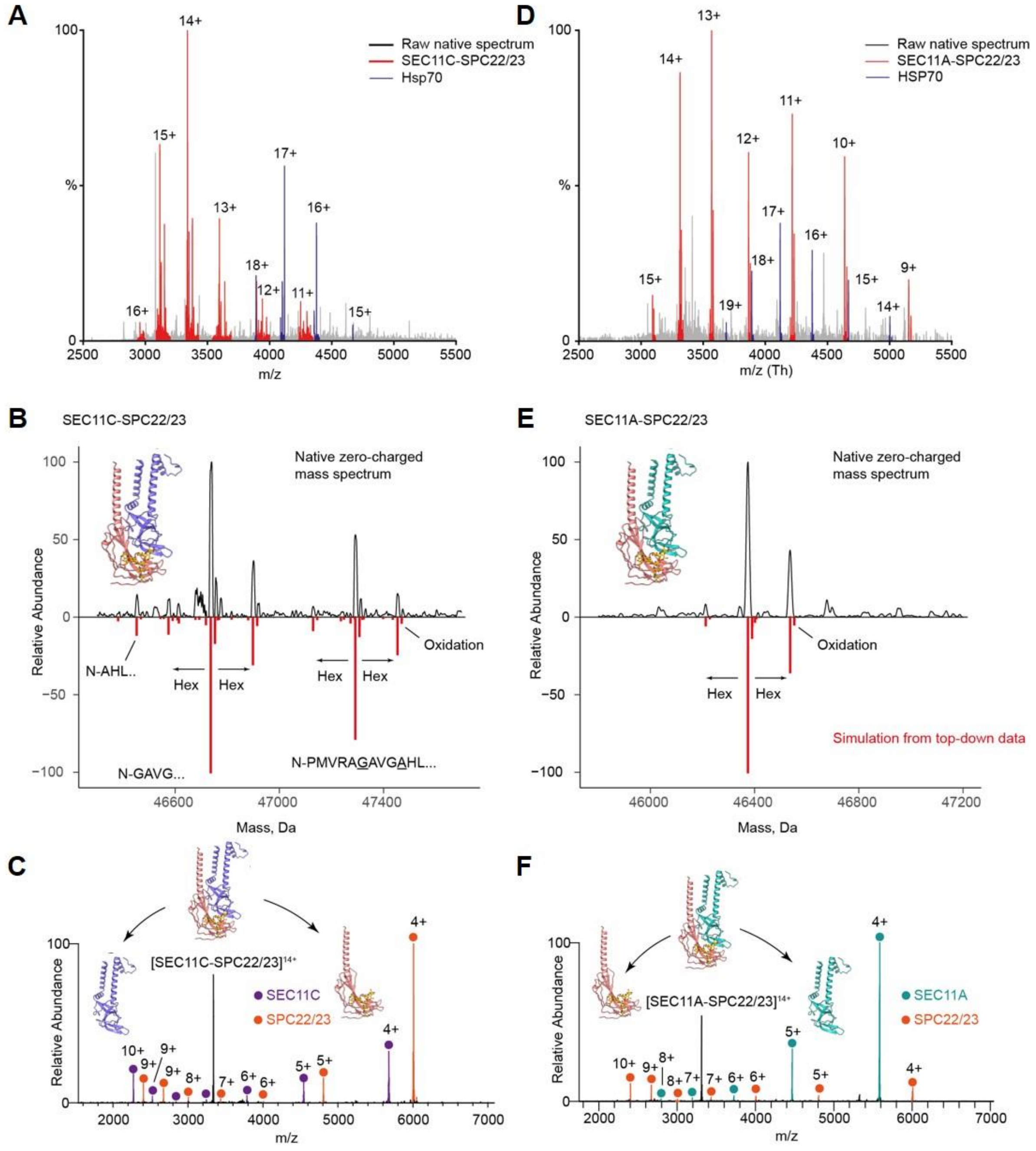


**Figure S8 | Native MS analysis of SPC22/23-SEC11 subcomplexes in SPC-A and SPC-C.** Samples were prepared in 0.15 M ammonium acetate containing 0.01% β-DDM. (**A**) Raw native MS spectrum demonstrating the SPC22/23-SEC11A dimer as the primary species in the SPC-A sample. The peripheral subunits SPC12 and SPC25 are likely lost when the samples are buffer exchanged. (**B**) Identification of SEC11C-SPC22/23 in the deconvoluted zero-charged spectrum by comparison with simulated mass spectrum from top-down MS. Primary PTMs contributing to microheterogeneity are annotated in the simulated spectrum. (**C**) Gas-phase activation of the most abundant 14+ charge state of SEC11C-SPC22/23 dimer confirms the oligomeric state and identity of non-covalently attached subunits. (**D**-**F**), Same as **A**-**C** but for the SPC-A complex.

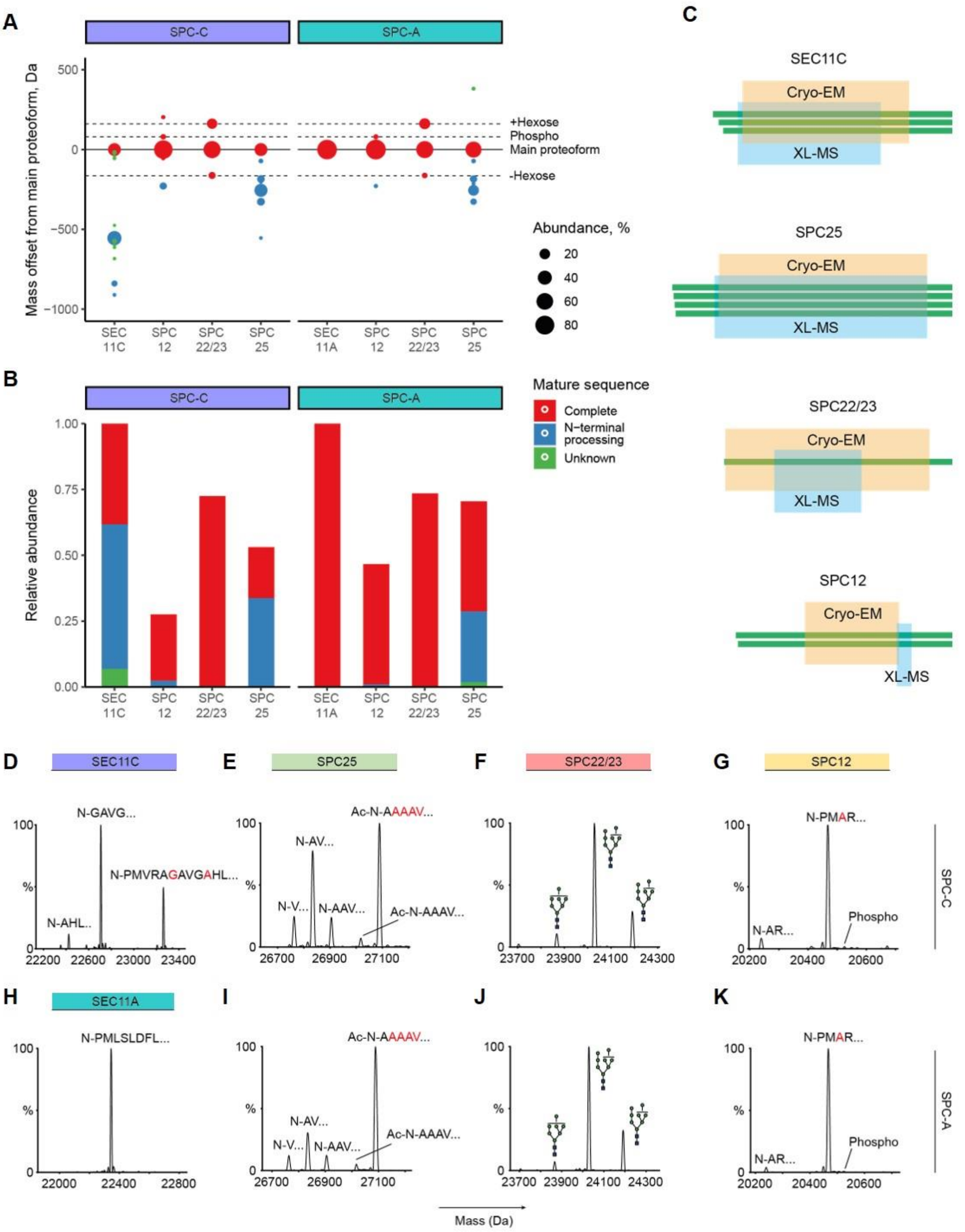


**Figure S9 | Top-down mass spectrometry reveals mature sequences and SPC proteoforms.** (**A**) Proteoform mass offsets determined for SPC subunits. Each offset is defined by the difference between a mass of a proteoform and the mass of the primary proteoform. The most prominent PTMs are highlighted with dashed lines. Main proteoforms are highlighted by a solid line. Apart from SEC11C and the N-glycosylated form of SPC22/23, the main proteoforms of all SPC subunits are unmodified. Circle size represents the fractional abundance of a proteoform per subunit. (**B**) Abundances of SPC subunits demonstrating the contribution of complete sequences, sequences with N-terminal processing, and undetermined sequence processing. (**C**) Sequences of distinct SPC proteoforms in the context of sequence coverages achieved with cryo-EM (orange) and XL-MS (blue). (**D-K)** Mass profiles displaying the primary proteoforms of SPC subunits for SPC-C (**D-G**) and SPC-A (**H-K**) complexes. For subunits with N-terminal sequence processing, a few N-terminal amino acids are displayed above the corresponding peaks. SPC25 was identified in N-acetylated and in N-terminally truncated, non-acetylated forms. For SPC22/23, the identified N-glycans are indicated above the corresponding proteoform peaks. A low-stoichiometric phosphorylated form of SPC12 is annotated in (**G**,**K**).

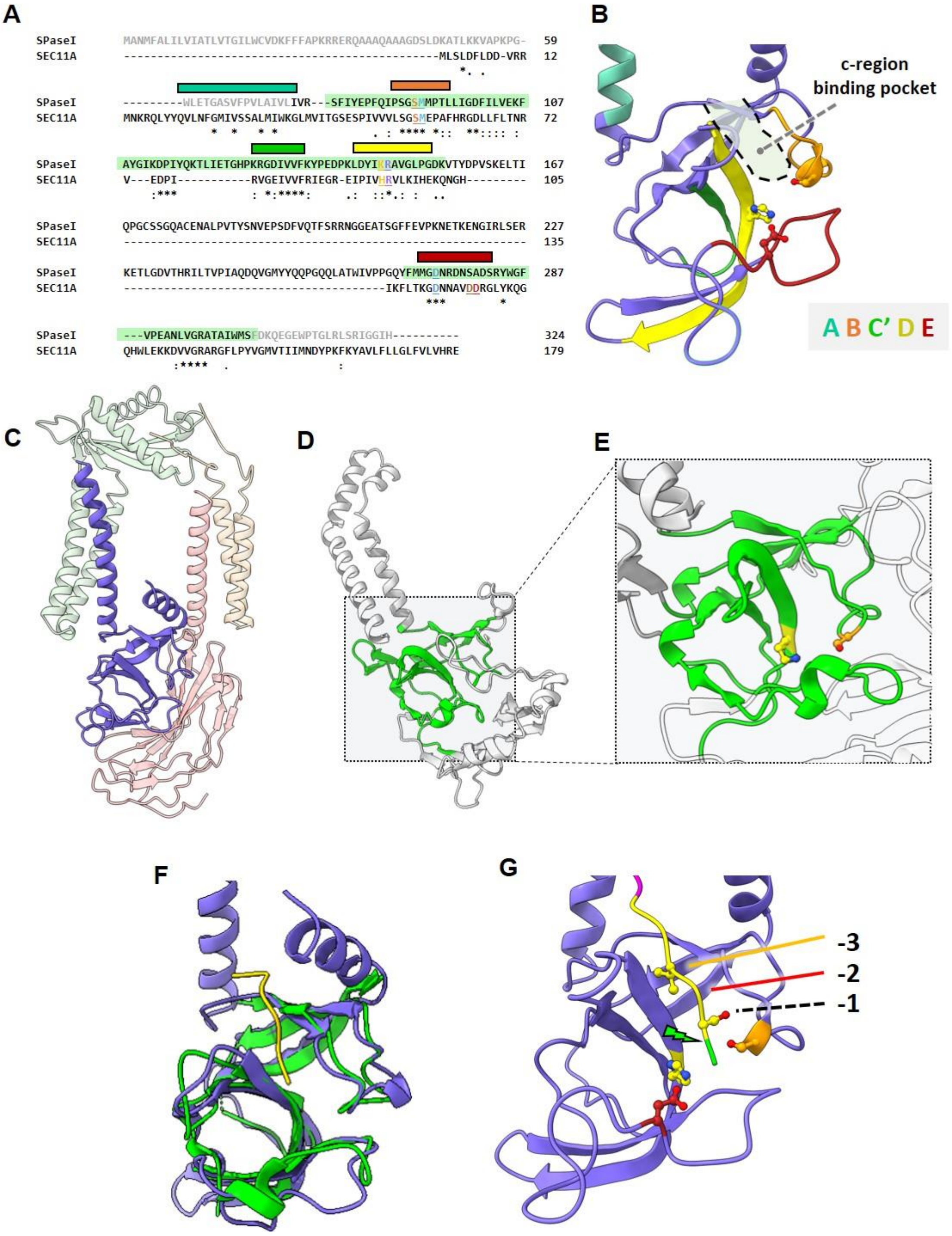


**Figure S10 | Comparison of human SEC11 and *E.coli* SPase I reveals the c-region binding pocket**. (**A**) Pairwise sequence alignment of SEC11A (UniProtKb ID P67812) and *E.coli* SPase I (UniProtKb ID P00803). Conserved boxes are highlighted by color (A = turquoise, B = orange; C' = green; D = yellow; E =brown). Sequence of the SPase I core highlighted in green. (**B**) The conserved boxes map to the transmembrane segment and SP binding pocket of SEC11. (**C**) Atomic model of SPC-C, colored as in Fig. 1. (**D**) trRosetta model of full-length *E.coli* SPase I. The catalytic domain is highlighted in green. (**E**) Zoom- in into the SP binding pocket of *E.coli* SPase I. The catalytic Ser-Lys dyad is highlighted. (**F**) Superposition of the luminal SEC11C portion with *E.coli* SPase I complexed with the lipopeptide inhibitor arylomycin (PDB-ID 1T7D, inhibitor in yellow). (**G**) Based on **F,** the c-region of bovine pre-prolactin was tentatively placed into the SEC11C binding pocket. In this hypothetic model, as in the bacterial enzyme, the inhibitor is forced into a beta strand conformation. The -1 and -3 positions point towards the shallow, hydrophobic pocket. The scissile bond (indicated with a green arrow) is located close to the catalytic residues.

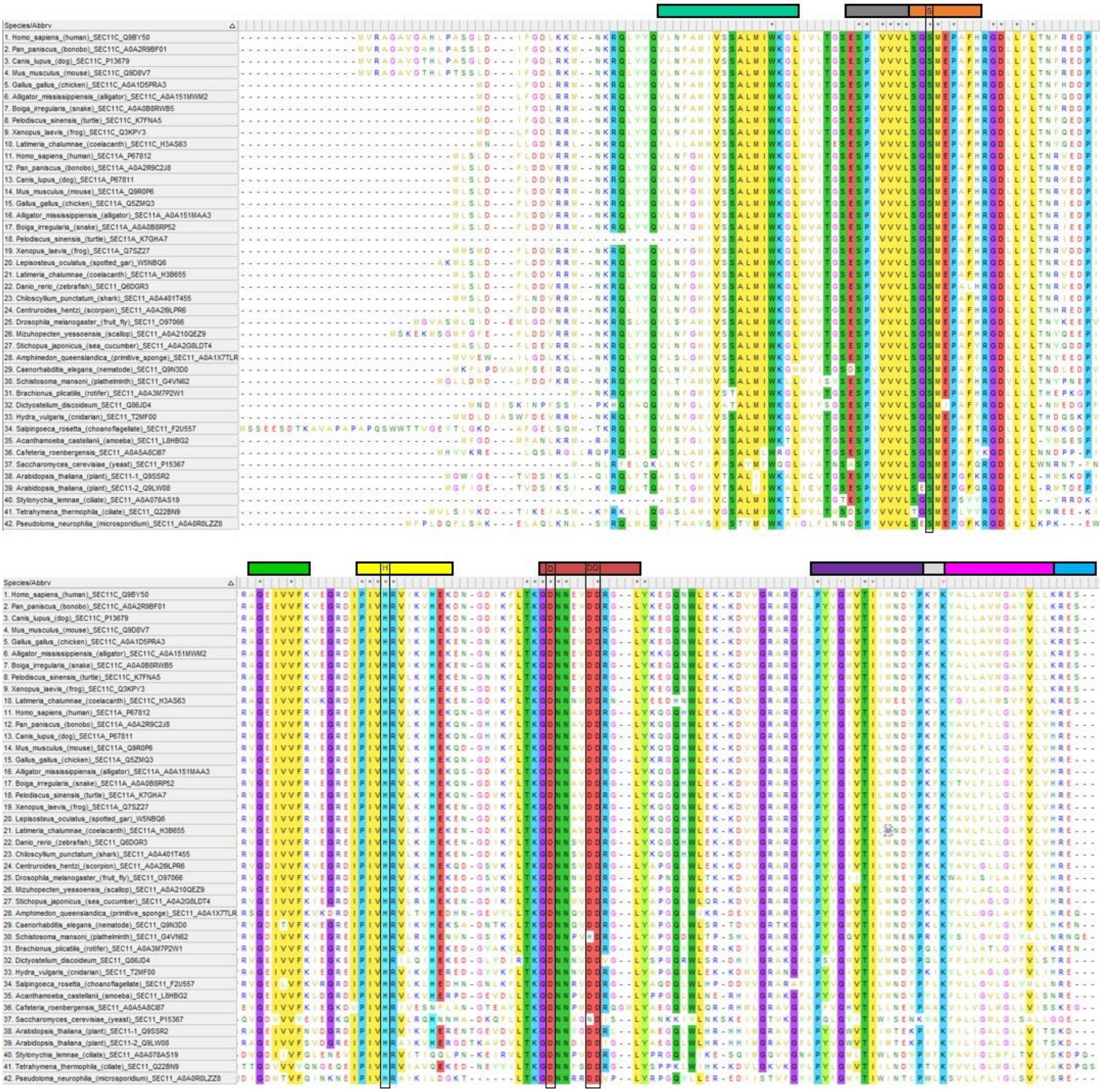

**Figure S11 | Residue conservation inSEC11 among eukaryotes.** Conserved boxes are highlighted as in Fig. S10. Catalytic Ser, His and the candidate Asp residues are highlighted in black boxes. Grey = additional region conserved in eukaryotes involved in binding pocket formation; purple = bowsprit helix, grey/magenta/cyan = segments of the pseudo-SP helix.

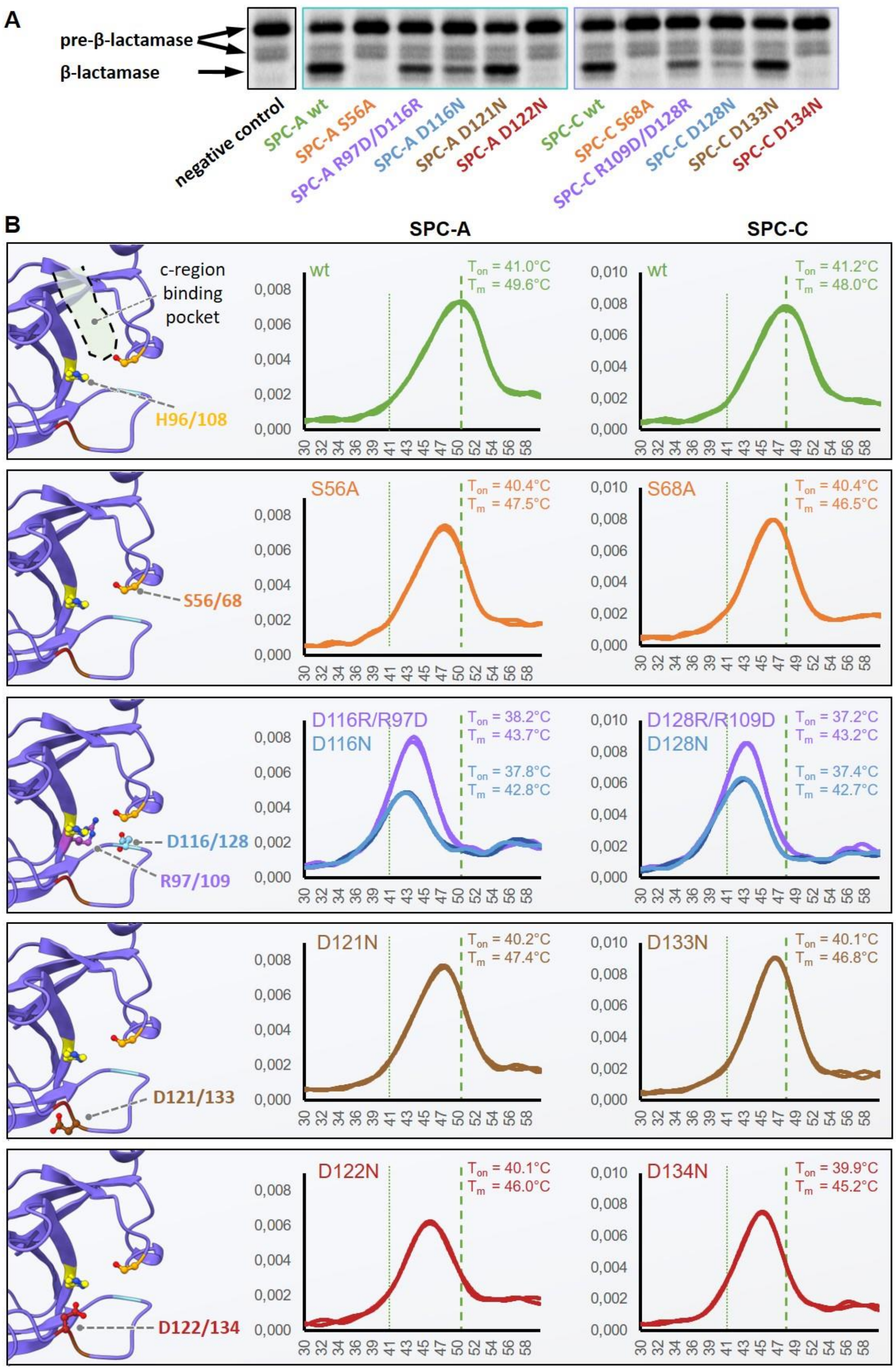


**Figure S12 | SPC is a serine protease with a catalytic triad.** (**A**) In vitro cleavage assay showing the processing of radiolabeled pre-β-lactamase in digitonin by SPC-A and SPC-C mutants. Negative control = no SPC added. (**B**) nano-differential scanning fluorimetry showing how the respective mutations affect protein stability. Left panel: Location of the respective residue (coloring as in Fig. 2). Middle and right panel: The first derivative of the fluorescence ratio at 350 nm and 330 nm (Y axis) is plotted against the temperature in °C (x axis). Melting onset ($T_{on}$) and melting temperature ($T_m$) are listed for each mutant. The melting profiles were determined as duplicates.

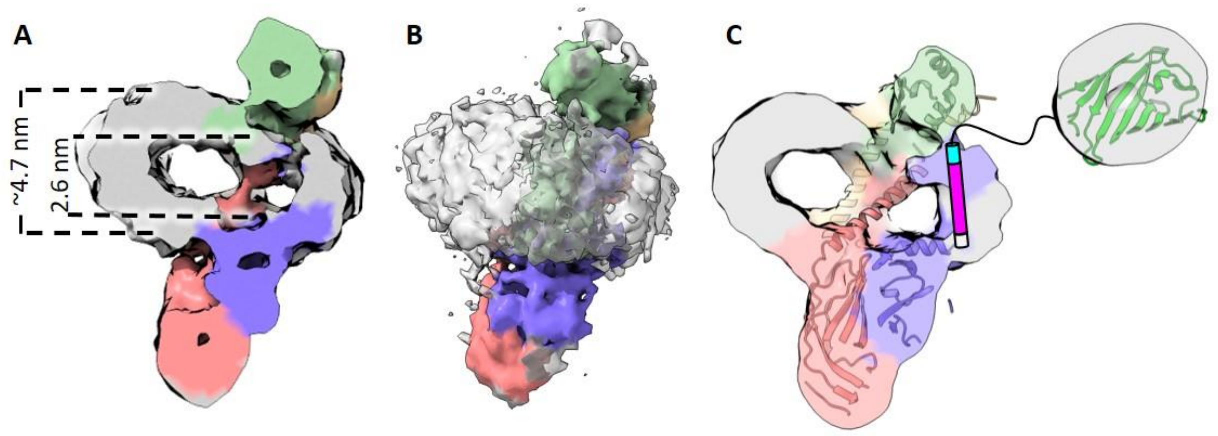


**Figure S13 | Membrane thinning in digitonin.** (**A**) Clipped view of SPC-C in digitonin at 12 Å resolution. The observed membrane thinning effect is akin to **Fig. 3A**. Coloring according to **Fig 3**, digitonin micelle depicted in grey. (**B**) Full view of SPC-C reconstituted in digitonin. (**C**) GFP (green cartoon) fused to the C-terminus of SEC11C is located at the cytosolic face of the particle, indicating that the C-terminus of SEC11C contains a transmembrane segment, termed here the pseudo-SP helix (indicated in grey/magenta/cyan).

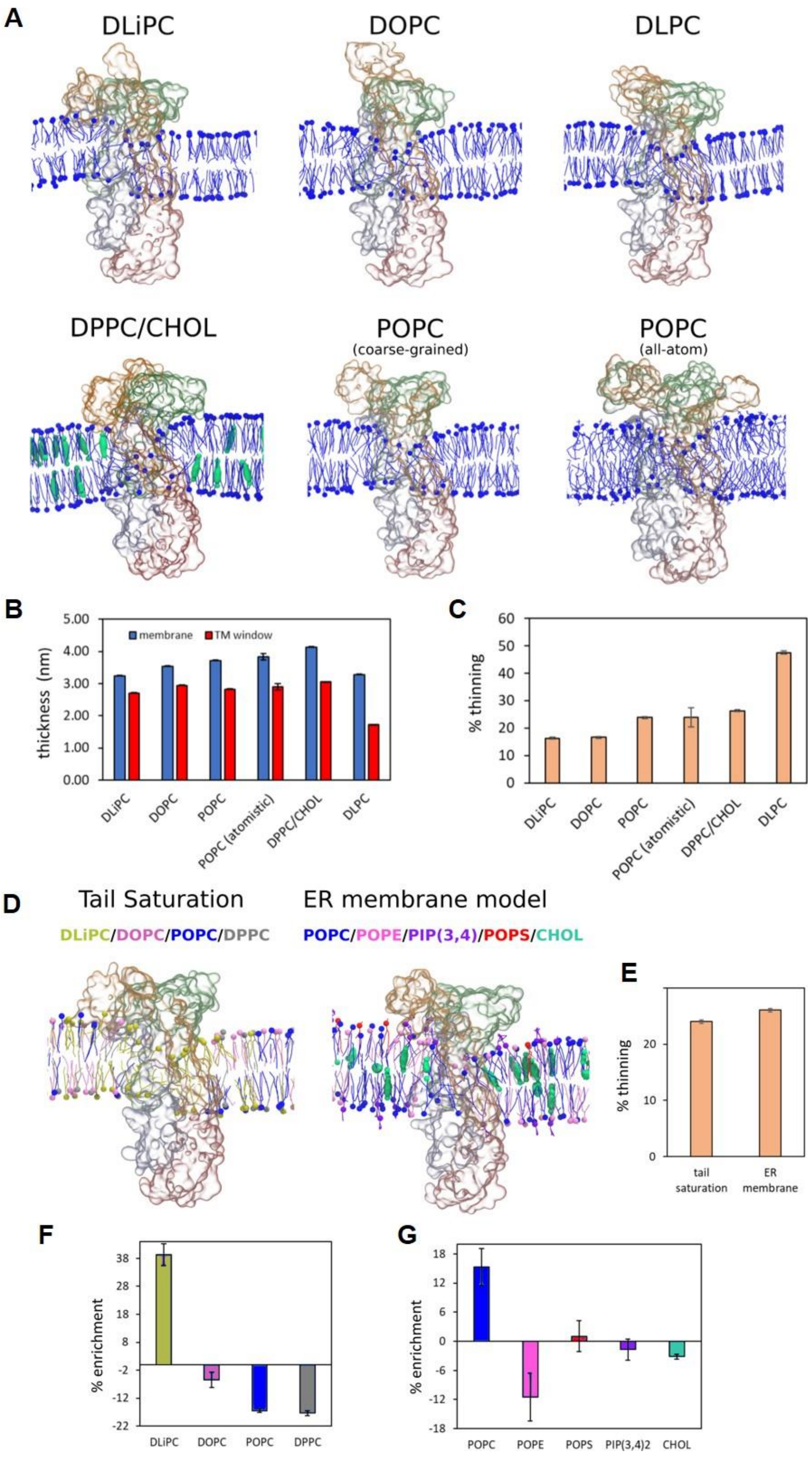


**Figure S14 | Lipid enrichment and membrane thinning induced by the TM window of SPC.** (**A**-**C**) Thinning induced in simple lipid bilayers. (**A**) Representative snapshots of coarse-grained MD simulations with SPC embedded in DLiPC, DOPC, DLPC, POPC and DPPC/CHOL (abbreviations see Table S3). For POPC, the results obtained with atomistic simulations are also shown. Coordinates were smoothed by averaging the coordinates of 4 neighbor frames of the trajectory. Only a slice of the bilayer around the TM window of SPC is highlighted. Blue = Phospholipids; cyan = cholesterol. SPC is represented by a transparent surface, with each protein chain identified by the same colors used in Fig. 3 of the main text. (**B**) Bilayer thickness of the bulk membrane (without SPC, blue) and in the TM window (red) obtained in the MD simulations. (**C**) Percentage of thinning induced by SPC in different membrane environments. (**D-G**) Thinning in complex membranes. (**D**) Representative snapshots of coarse-grained MD simulations of SPC embedded in two complex membrane environment: (i) in the left panel, a mixture of PC lipids with different level of tail saturations; (ii) in the right panel, an endoplasmic reticulum (ER) membrane model. For more details about lipid ratio, see **Table S3**. (**E**) Percentage of thinning induced by SPC in the complex lipid mixtures. (**F**) and (**G**) Percentage of lipid enrichment in the TM window region in relation to the lipid ratio of bulk membrane. Negative values indicate depletion of lipids. Results are displayed for the mixture of PC lipids with different level of tail saturations (**F**) and for the ER membrane model (**G**).

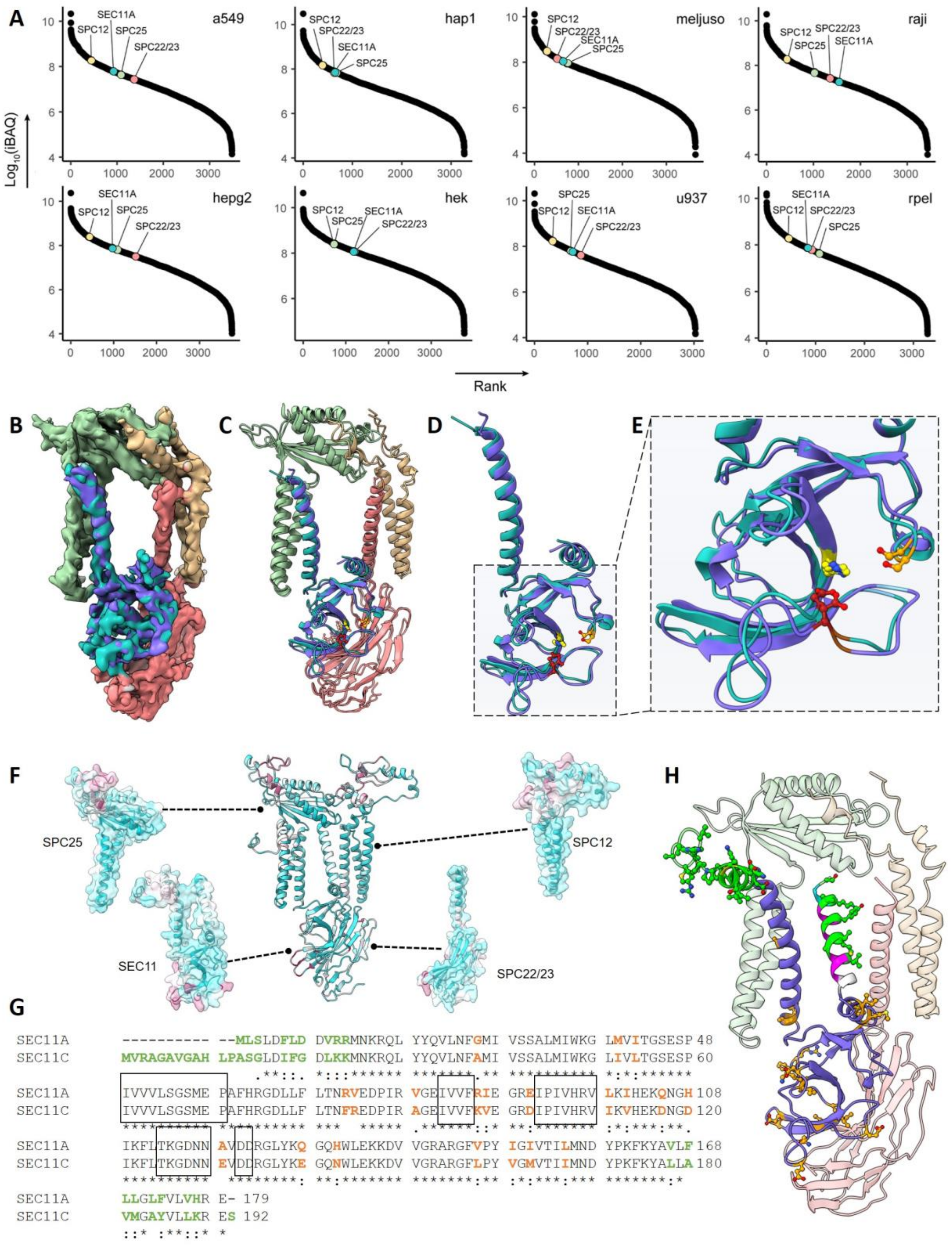


**Figure S15 | Differences between SEC11A and SEC11C**. (**A**) iBAQ intensities of SPC proteins determined for eight different cell lines are plotted by rank. SPC12, SPC25, SPC22/23, and SEC11A are marked; SEC11C was not detected in any of the tested cell lines. (**B**) Superposition of density maps for SPC-A and SPC-C (colored according to Fig. 1). (**C**) Superposition of atomic models for SPC-A and SPC-C. (**D**-**E**)**,** Superposition of atomic models for SEC11A (teal) and SEC11C (purple). (**F**) Conservation mapped onto the SPC structure (teal = conserved, red = variable). Contact points between subunits as well as the SP binding groove are highly conserved. (**G**) Pairwise alignment of SEC11A and SEC11C. Residues that form the SP binding site are highlighted in black boxes. Variable residues that are resolved in the maps are highlighted in orange, variable residues in flexible regions are highlighted in green. (**H**) differences between SEC11A and SEC11C mapped onto the SPC-C structure, colored as in **F**.

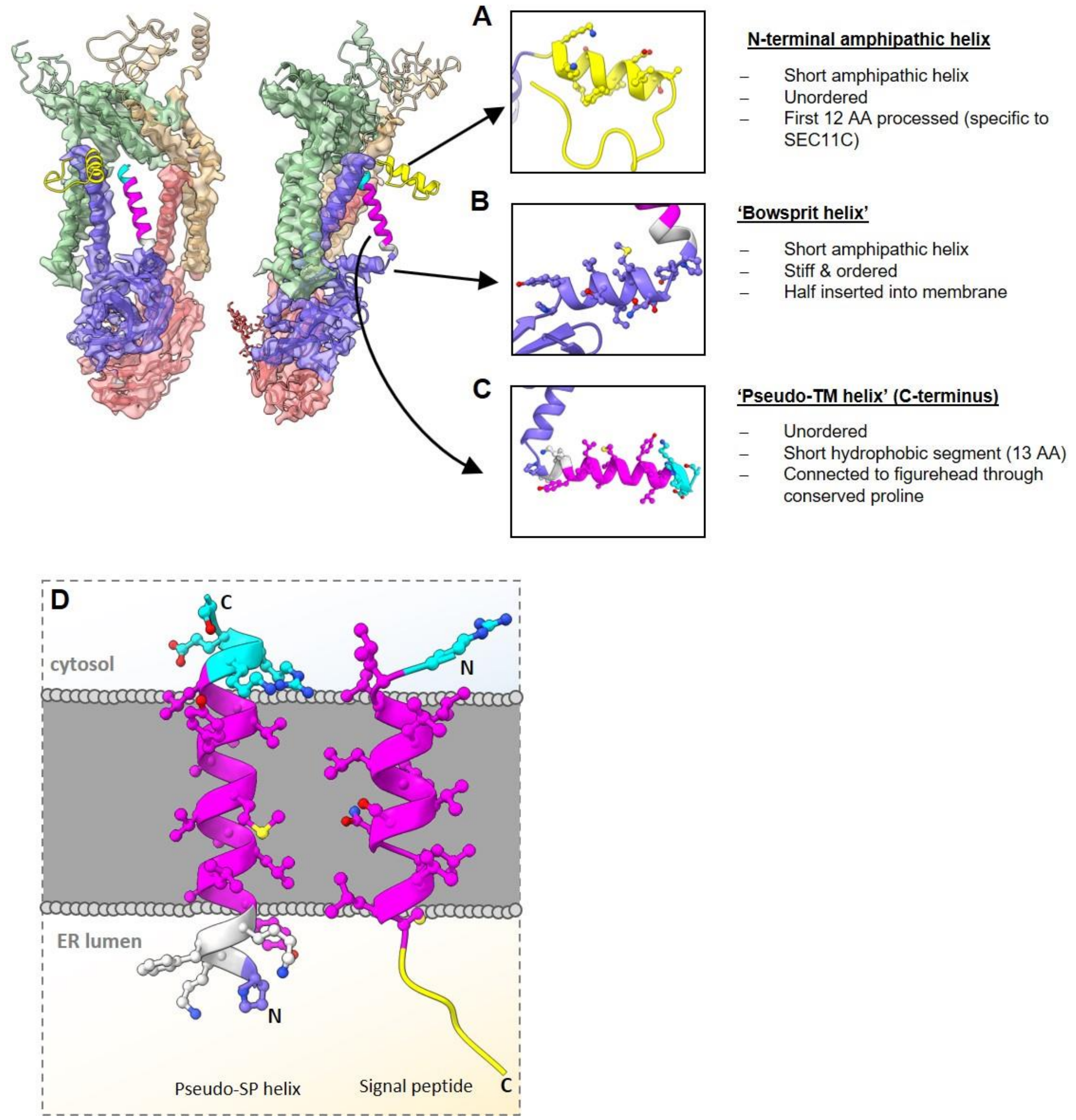


**Figure S16 | Unresolved regions of SEC11A/C.** Unresolved regions of the SPC are shown as predicted by trRosetta. (**A-C**) Zoom-in and characteristics of terminal SEC11 helices. (**D**) comparison of the pseudo-SP helix of SEC11C to pre-prolactin SP (PDB ID 3JC2).

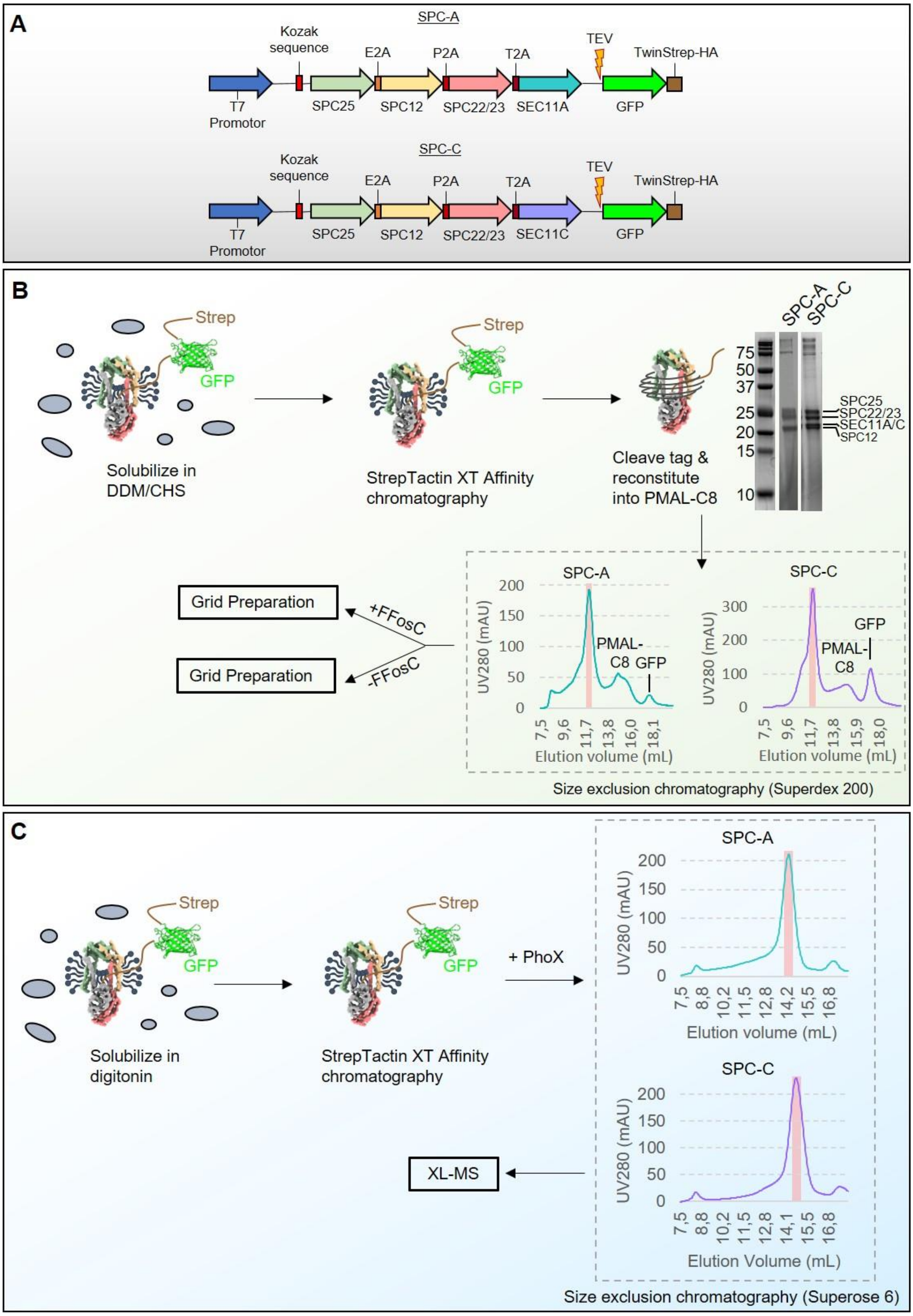


**Figure S17 | Purification of human SPC for cryo-EM and XL-MS.** (**A**) Expression constructs used for SPC purification. (**B**) Purification for cryo-EM. FFosC = fluorinated Fos-choline. (**C**) Purification for XL-MS. Workflows were analogous for SPC-A (teal) and SPC-C (purple). SEC11A/C are indicated in grey in panels B and C**,** the other subunits are colored according to a. Monodisperse SEC peaks used for analysis are highlighted in red. PhoX = crosslinker.

### Supplementary Tables

**Table S1 | Data collection and refinement statistics**

| | SPC-A | SPC-C |
|---|---|---|
| **Data collection and processing** | | |
| **Microscope** | Talos Arctica | Talos Arctica |
| **Camera** | K2 summit | K2 summit |
| **Magnification** | 165,000 | 165,000 |
| **Voltage (kV)** | 200 | 200 |
| **Electron exposure ($e^-$/Å$^2$)** | 60 | 60 |
| **Defocus range** | 0.5-4.0 | 0.5-4.0 |
| **Pixel Spacing (Å)** | 0.81 | 0.81 |
| **Symmetry imposed** | C1 | C1 |
| **Final Nr. particle images** | 29,508 | 60,598 |
| **Map Resolution** | 4.9 | 4.9 |
| FSC Threshold | 0.143 | 0.143 |
| **Map sharpening B factor (Å$^2$)** | -180 | -180 |
| **Model validation** | | |
| **MolProbity Score** | 1.7 | 1.5 |
| **Clashscore** | 4.5 | 3.9 |
| **Rotamer Outliers (%)** | 0.2 | 0.0 |
| **Ramachandran plot** | | |
| Favored | 91.9 | 95.0 |
| Allowed | 100.0 | 100.0 |
| Outliers | 0.0 | 0.0 |
| **Real-space correlation** | 0.68 | 0.73 |
| **Mean model B factor (Å$^2$)** | 144 | 231 |

**Table S2 | Mass Spectrometry data availability**

| | Associated files | Experiment type | Pa-nels | Comments |
|---|---|---|---|---|
| Fig. S7 | 20191026_F1_Ag5_Steig002_SA_SPC_C12_KonstrA_E (.raw/ .txt)<br>20191026_F1_Ag5_Steig002_SA_SPC_D12_KonstrA_E (.raw / .txt)<br>20191030_F1_Ag5_Steig002_SA_SPC_C11_KonstrB_E (.raw / .txt)<br>20191030_F1_Ag5_Steig002_SA_SPC_C12D12_KonstrB_E(.raw/.txt) | XL-MS | all | Crosslinks on SPC complex after SEC purification |
| Fig. S15 | 20200714_OR16_UM7_Tamar002_SA_HEK_TRYPSIN_115min.raw<br>20200714_OR16_UM7_Tamar002_SA_HEK_TRYPSIN_115min_2.raw<br>20200714_OR16_UM7_Tamar002_SA_HEK_TRYPSIN_115min_3.raw<br>20200723_OR16_UM7_Tamar002_SA_A549_115min_1.raw<br>20200723_OR16_UM7_Tamar002_SA_A549_115min_2.raw<br>20200723_OR16_UM7_Tamar002_SA_A549_115min_3.raw<br>20200723_OR16_UM7_Tamar002_SA_HAP1_115min_1.raw<br>20200723_OR16_UM7_Tamar002_SA_HAP1_115min_2.raw<br>20200723_OR16_UM7_Tamar002_SA_HAP1_115min_3.raw<br>20200723_OR16_UM7_Tamar002_SA_HEPG2_115min_1.raw<br>20200723_OR16_UM7_Tamar002_SA_HEPG2_115min_2.raw<br>20200723_OR16_UM7_Tamar002_SA_HEPG2_115min_3.raw<br>20200723_OR16_UM7_Tamar002_SA_MELJUSO_115min_1.raw<br>20200723_OR16_UM7_Tamar002_SA_MELJUSO_115min_2.raw<br>20200723_OR16_UM7_Tamar002_SA_MELJUSO_115min_3.raw<br>20200723_OR16_UM7_Tamar002_SA_RAJI_115min_1.raw<br>20200723_OR16_UM7_Tamar002_SA_RAJI_115min_2.raw<br>20200723_OR16_UM7_Tamar002_SA_RAJI_115min_3.raw<br>20200723_OR16_UM7_Tamar002_SA_RPEL_115min_1.raw<br>20200723_OR16_UM7_Tamar002_SA_RPEL_115min_2.raw<br>20200723_OR16_UM7_Tamar002_SA_RPEL_115min_3.raw<br>20200723_OR16_UM7_Tamar002_SA_U937_115min_1.raw<br>20200723_OR16_UM7_Tamar002_SA_U937_115min_2.raw<br>20200723_OR16_UM7_Tamar002_SA_U937_115min_3.raw<br>evidence.txt | Shot-gun proteomics | a | SPC subunits detected in different cell lines with bottom-up LC-MS/MS |
| Fig. S8 | Tamar002_UHMR_spcA.raw<br>Tamar002_UHMR_spcA_HCD125.raw<br>Tamar002_UHMR_spcA_HCD150.raw<br>Tamar002_UHMR_spcC.raw<br>Tamar002_UHMR_spcC_HCD125.raw<br>Tamar002_UHMR_spcC_HCD150.raw | Native MS | all | Native MS data for SPC-A and SPC-C, and HCD spectra of z=14+ of SPC22/23-SEC11A and SPC22/23-SEC11C. |
| Fig. 1 | Tamar002_SA_SPC_A_STREP_LOHI_ETD.raw<br>Tamar002_SA_SPC_A_STREP_LOHI_ETD.xlsx<br>Tamar002_SA_SPC_A_STREP_LO_FMS.csv<br>Tamar002_SA_SPC_A_STREP_LO_FMS.raw<br>Tamar002_SA_SPC_C_FLAG_LOHI_ETD.raw<br>Tamar002_SA_SPC_C_FLAG_LOHI_ETD.xlsx<br>Tamar002_SA_SPC_C_FLAG_LO_FMS.csv<br>Tamar002_SA_SPC_C_FLAG_LO_FMS.raw | TD-MS | b, c | Quantification of SPC proteins in FLAG and STREP pull-downs |
| Fig. S9 | Tamar002_SA_SPC_A_LOHI_ETD.raw<br>Tamar002_SA_SPC_A_LOHI_ETD.xlsx<br>Tamar002_SA_SPC_A_LO_FMS.csv<br>Tamar002_SA_SPC_A_LO_FMS.raw<br>Tamar002_SA_SPC_C_LOHI_ETD.raw<br>Tamar002_SA_SPC_C_LOHI_ETD.xlsx<br>Tamar002_SA_SPC_C_LO_FMS.csv<br>Tamar002_SA_SPC_C_LO_FMS.raw | TD-MS | a-k | Overview of proteoforms detected in top-down and intact mass MS of SPC-A and SPC-C complexes |

**Table S3 | Summary of the MD simulations performed in this study.** All simulations were performed with and without SPC embedded in the membranes.

| Resolution | Lipid Composition* | Duration |
|---|---|---|
| **Coarse-Grained** (**Martini 3**) | DLPC | 20 µs |
| | DLiPC | 20 µs |
| | DOPC | 20 µs |
| | POPC | 20 µs |
| | DPPC, CHOL (0.70:0.30) | 20 µs |
| | DLiPC, DOPC, POPC, DPPC (0.25: 0.25: 0.25: 0.25) | 20 µs |
| | Endoplasmic reticulum membrane model POPC, POPE, POPS, PI(3,4)P2, CHOL (0.442:0.255:0.0425:0.1105:0:15) | 20 µs |
| **All-Atom** (**CHARMM 36m**) | POPC | 2x 150 ns |

*Abbreviations: DLPC=1,2-dilauroyl-sn-glycero-3-phosphocholine; DLiPC=1,2-dilinoleoyl-sn-glycero-3-phosphocholine; DOPC=1,2-dioleoyl-sn-glycero-3-phosphocholine; POPC=-palmitoyl-2-oleoyl-sn-glycero-3-phosphocholine; DPPC=1,2-dipalmitoyl-sn-glycero-3-phosphocholine; CHOL=cholesterol; POPE= 1-Palmitoyl-2-oleoyl-sn-glycero-3-phosphoethanolamine; POPS=1-palmitoyl-2-oleoyl-sn-glycero-3-phospho-L-serine; PI(3,4)P2=phosphatidylinositol 3,4-bisphosphate

### Supplementary Movies

**Movie S1 | Structures of SPC-A and SPC-C.** The movie compares the density maps obtained for SPC-C and SPC-A (colored as in **Fig. 2**) and depicts the map-to-model fit of both atomic models. The models are superposed, and the catalytic residues are highlighted as ball-and-stick models. N-glycosylation at Asp141 of SPC22/23 is also depicted, although not visible at the given threshold.

**Movie S2 | Membrane thinning by SPC-C.** The map-to model fit of SPC-C is shown (colored as in **Fig. 2**). The signal peptide of bovine pre-prolactin (represented as ribbons, colored as in **Fig. 4A**) was placed into the putative SP binding pocket using *Coot*. The -1 and -3 positions of the SP c-region are highlighted as ball-and-stick models as in **Fig. S10G**. The Gaussian-filtered, experimentally determined density for the PMAL-C8 micelle is shown at different threshold levels in order to depict the membrane thinning within the TM window. The thickness of the micelle in the putative h-region binding pocket is consistent with the length of the SP h-region.

**Movie S3 | A glimpse of the membrane thinning induced by SPC-C in an ER membrane model during 2μs of a coarse-grained MD simulation.** Coordinates were smoothed by averaging the coordinates of 4 neighbor frames of the trajectory. Only a slice of the membrane around the TM window of SPC is highlighted. Lipids are shown with different colors: POPC (blue), POPS (red), POPE (pink), PI(3,4)2 (purple) and cholesterol (green). SPC-C is represented by a transparent surface, with each protein chain identified by the same colors used in **Fig. 3**.
